# XDH-1 inactivation causes xanthine stone formation in *C. elegans* which is inhibited by SULP-4-mediated anion exchange in the excretory cell

**DOI:** 10.1101/2025.01.24.634556

**Authors:** Jennifer Snoozy, Sushila Bhattacharya, Brandon Johnson, Robin R. Fettig, Ashley Van Asma, Chloe Brede, Sophia G. Miller, Martina Ralle, Kurt Warnhoff

**Affiliations:** Pediatrics and Rare Diseases Group, Sanford Research, Sioux Falls, SD 57104, USA; Department of Pediatrics, Sanford School of Medicine, University of South Dakota, Sioux Falls, SD 57105 USA; Department of Basic Biomedical Sciences, Sanford School of Medicine, University of South Dakota, Vermillion, SD 57069, USA; Department of Molecular and Medical Genetics, Oregon Health & Science University, Portland, OR 97201, USA

## Abstract

Xanthine dehydrogenase (XDH) is a molybdenum cofactor (Moco) requiring enzyme that catabolizes hypoxanthine into xanthine and xanthine into uric acid, the final steps in purine catabolism. Human patients with mutations in XDH develop xanthinuria which can lead to xanthine stones in the kidney, recurrent urinary tract infections, and renal failure. Currently there are no therapies for treating human XDH deficiency. Thus, understanding mechanisms that maintain purine homeostasis is an important goal of human health. Here, we used the nematode *C. elegans* to model human XDH deficiency using 2 clinically relevant paradigms: Moco deficiency or loss-of-function mutations in *xdh-1,* the *C. elegans* ortholog of XDH. Both Moco deficiency and *xdh-1* loss of function caused the formation of autofluorescent xanthine stones in *C. elegans*. Surprisingly, only 2% of *xdh-1* null mutant *C. elegans* developed a xanthine stone, suggesting additional pathways may regulate this process. To uncover such pathways, we performed a forward genetic screen for mutations that enhance the penetrance of xanthine stone formation in *xdh-1* null mutant *C. elegans*. We isolated multiple loss-of-function mutations in the gene *sulp-4* which encodes a sulfate permease homologous to human SLC26 anion exchange proteins. We demonstrated that SULP-4 acts cell-nonautonomously in the excretory cell to limit xanthine stone accumulation. Interestingly, *sulp-4* mutant phenotypes were suppressed by mutations in genes that encode for cystathionase (*cth-2)* or cysteine dioxygenase (*cdo-1*), members of the sulfur amino acid catabolism pathway required for production of sulfate, a substrate of SULP-4. We propose that sulfate accumulation caused by *sulp-4* loss of function promotes xanthine stone accumulation. We speculate that sulfate accumulation causes osmotic imbalance, creating conditions in the intestinal lumen that favor xanthine stone accumulation. Supporting this model, a mutation in *osm-8* that constitutively activates the osmotic stress response also promoted xanthine stone accumulation in an *xdh-1* mutant background. Thus, our work establishes a *C. elegans* model for human XDH deficiency and identifies the sulfate permease *sulp-4* as a critical player controlling xanthine stone accumulation.

## Background

Purines are an abundant and fundamental metabolite class that are essential for the generation of RNA and DNA molecules and purine nucleotides are critical energy sources (ATP) and signaling molecules (GTP). Failures in purine metabolism can lead to both common and rare diseases. For instance, oncogenic mutations activate nucleotide biosynthetic capacity in diverse cancers, promoting cancer progression [1]. Mutations in enzymes in the purine metabolic pathway cause rare inborn errors of metabolism such as Lesch-Nyhan syndrome, purine nucleoside phosphorylase deficiency, and xanthinuria [2–5]. Thus, understanding the mechanisms that impact purine homeostasis is an important goal of human health.

Xanthinuria, an inborn error of purine metabolism, is caused by inactivation of xanthine dehydrogenase (XDH), the terminal enzyme in purine catabolism that oxidizes hypoxanthine to xanthine and xanthine to uric acid [6] (**Fig. 1A**). There are two types of human xanthinuria; type I is caused by mutations in the gene encoding the xanthine dehydrogenase enzyme and type II is caused by mutations in genes essential for the synthesis of the molybdenum cofactor, an essential prosthetic group for XDH [4, 5]. Both forms of xanthinuria present with high levels of xanthine in the urine and low levels of uric acid which can result in the formation of xanthine stones in kidneys and muscles, sometimes causing renal failure. There is currently no curative therapy for xanthinuria, however high fluid intake and low purine diets are recommended for patients [7].

**Figure 1:**
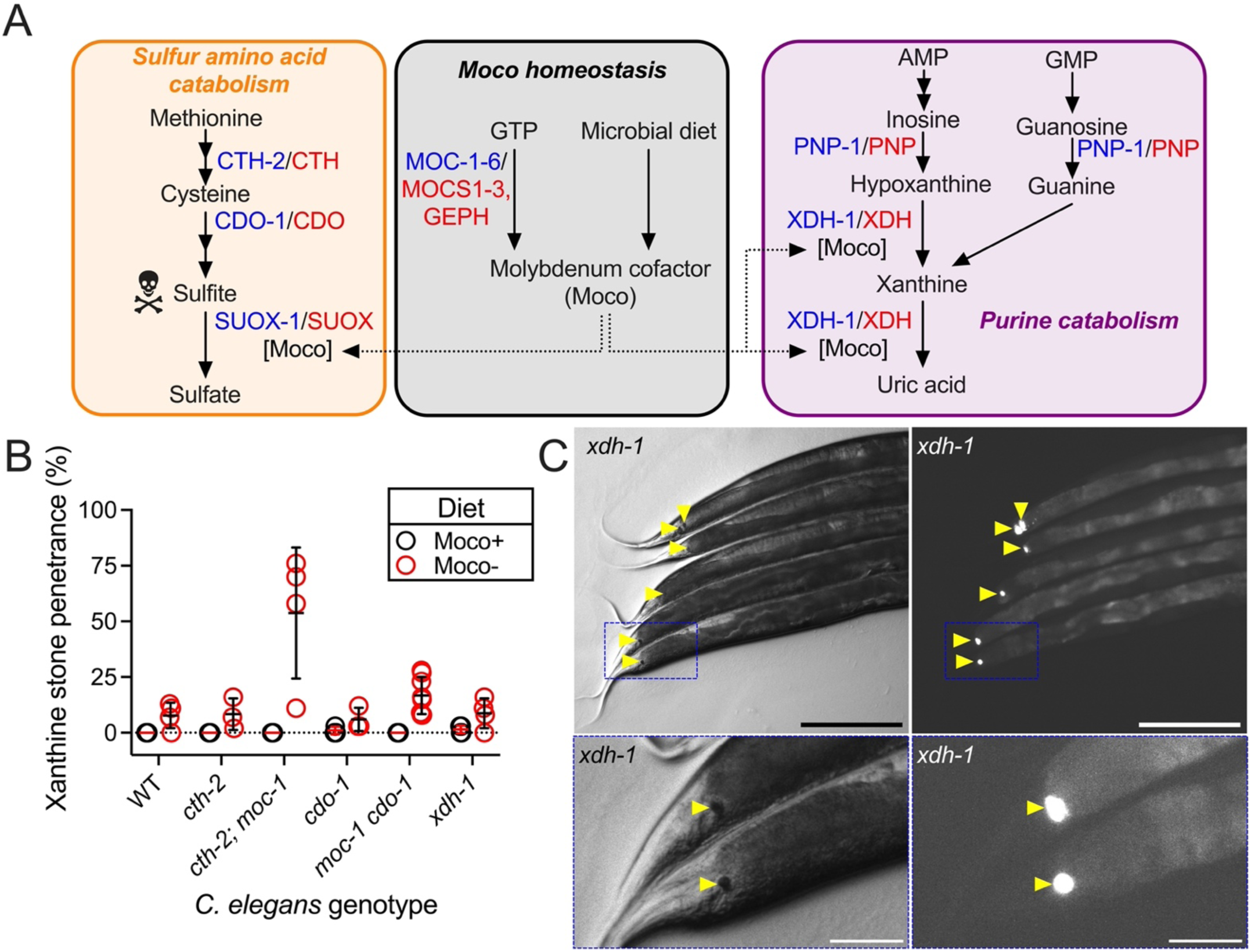
Moco deficiency or loss of *xdh-1* promoted the formation of autofluorescent xanthine stones. A) The role of Moco in *C. elegans* metabolism. We highlight i) pathways for Moco synthesis and import, ii) sulfur amino acid catabolism governed by cystathionase (CTH-2/CTH), cysteine dioxygenase (CDO-1/CDO), and Moco-requiring sulfite oxidase (SUOX-1/SUOX), and iii) purine catabolism controlled by purine nucleoside phosphorylase (PNP-1/PNP) and the Moco-requiring xanthine dehydrogenase (XDH-1/XDH). *C. elegans* enzymes (blue) and their human homologs (red) are displayed. B) Wild-type and mutant *C. elegans* were cultured on wild-type (black, Moco+) or *τιmoaA* mutant (red, Moco-) *E. coli* and assessed for the formation of xanthine stones over the first 3 days of adulthood. Individual data points represent biological replicates. Mean and standard deviation are displayed. Complete information regarding sample size and individuals scored per biological replicate are found in **Table S2**. C) Brightfield (left) and fluorescent (right) images of the posterior of *xdh-1(ok3234)* mutant adult *C. elegans* cultured on wild-type *E. coli.* Xanthine stones are highlighted (yellow arrowheads). The blue box indicates the region magnified in the lower panels. Scale bar is 250μm (top) or 50μm (bottom).

Molybdenum cofactor (Moco) is an essential prosthetic group required for development in animals ranging from the nematode *C. elegans* to humans [4, 8]. Moco is synthesized by an ancient and conserved biosynthetic pathway [9]. *C. elegans* has recently emerged as a powerful model system for studying Moco biology, and the genes that encode the Moco biosynthetic enzymes are termed *moc* in *C. elegans* for MOlybdenum Cofactor biosynthesis [8, 10]. In addition to endogenous Moco synthesis, *C. elegans* can also acquire and use Moco from its bacterial diet [8, 11, 12]. Given its genetic tractability and the ability to manipulate animal Moco content by simple dietary manipulation, *C. elegans* is a useful model for understanding the biology of Moco and Moco-requiring enzymes, such as XDH-1 (**Fig. 1A**).

Here, we genetically explored the formation of xanthine stones in *C. elegans,* which emerged during Moco deficiency or in *xdh-1* mutant *C. elegans,* mirroring type I and type II human xanthinuria [4, 5]. Surprisingly, only a small percentage of Moco-deficient and *xdh-1* mutant *C. elegans* developed xanthine stones, suggesting additional parallel pathways for limiting stone formation. To identify novel regulators of xanthine stone accumulation, we performed an unbiased chemical mutagenesis screen for mutations that enhanced the penetrance of xanthine stone formation in *xdh-1* mutant animals. In this screen we recovered five loss-of-function alleles of *sulp-4,* a gene which encodes a member of the sulfate permease family of transporters with homology to human SLC26 transporters [13]. We demonstrated that SULP-4 acts in the *C. elegans* excretory cell, analogous to the human kidney, to inhibit the accumulation of xanthine stones [14]. We further showed that *sulp-4* was required for normal development. Interestingly, we found that phenotypes caused by *sulp-4* loss of function were suppressed by inactivating mutations in *cth-2* or *cdo-1,* genes that encode core members of the sulfur amino acid catabolism pathway (**Fig. 1A**) [8]. We suggest that *sulp-4* loss of function causes sulfate accumulation. Disruption of *cth-2* or *cdo-1* impairs endogenous sulfate production, thereby suppressing the phenotypes observed in *sulp-4* mutant animals. Sulfate accumulation may cause an osmotic imbalance that leads to an increased rate of xanthine stone formation in the intestinal lumen. This model is supported by our observation that a mutation in *osm-8*, a key regulator of osmotic homeostasis, also promoted xanthine stone accumulation in an *xdh-1* mutant background [15]. Thus, our work establishes a *C. elegans* model for the rare genetic disease xanthinuria and identifies *sulp-4* as a potent genetic modifier of this disease pathology in *C. elegans,* likely acting through disrupted osmotic homeostasis.

## Results

### XDH-1 inactivation caused the accumulation of xanthine stones in *C. elegans*

To explore the pathology of Moco deficiency in the nematode *C. elegans,* we cultured *cth-2; moc-1* and *moc-1 cdo-1* double mutant animals on wild type (Moco replete) or *ΔmoaA* mutant (Moco deficient) *E. coli*. The *moc-1* mutation prevents the endogenous synthesis of Moco while the *cth-2* and *cdo-1* mutations suppress the lethality typically associated with animal Moco deficiency [8]. *cth-2* encodes cystathionase which converts cystathionine to cysteine and *cdo-1* encodes cysteine dioxygenase which oxidizes cysteine to cysteinesulfinate ultimately producing sulfites (**Fig. 1A**). Sulfites are extremely reactive and are detoxified to sulfate by the Moco-requiring sulfite oxidase (SUOX-1) enzyme. Thus, by changing the dietary *E. coli*, we can control whether the animals have Moco. When culturing *cth-2; moc-1* double mutant animals on Moco-*E. coli* we surprisingly observed animals that developed autofluorescent stones, typically found in the posterior of the intestine. 54% of *cth-2; moc-1* animals fed a Moco-diet developed an autofluorescent stone while 0% of *cth-2; moc-1* animals developed an autofluorescent stone when fed a diet that provided Moco. We observed similar results for *moc-1 cdo-1* double mutant animals where 17% of animals developed an autofluorescent stone on Moco-*E. coli* and 0% developed a stone when fed wild-type *E. coli* (**Fig. 1B**). Thus, we conclude that the formation of these autofluorescent stones is caused by Moco deficiency.

Surprisingly, we also observed the formation of autofluorescent stones in 8% of wild-type *C. elegans* when cultured on Moco-*E. coli.* 0% of wild-type animals developed an autofluorescent stone when fed Moco+ *E. coli* (**Fig. 1B**). This result demonstrates that dietary Moco deficiency alone is sufficient to promote the formation of autofluorescent stones. This result is surprising as wild-type animals are still competent to produce Moco through their endogenous biosynthetic pathway. However, these data are consistent with our recent findings that the *C. elegans* diet plays a significant role in promoting Moco homeostasis [12].

Given that the development of these autofluorescent stones was dependent upon dietary Moco deficiency, we reasoned that the phenotype was likely being caused by inactivation of one of the four animal Moco-requiring enzymes (sulfite oxidase, xanthine dehydrogenase, aldehyde oxidase, and mitochondrial amidoxime reducing component) [9]. Interestingly, inactivation of the Moco-requiring enzyme xanthine dehydrogenase causes the accumulation of insoluble and fluorescent xanthine stones in organisms as diverse as plants, fruit flies, and humans [5, 16–20]. We therefore hypothesized that the autofluorescent stones we observed during *C. elegans* Moco deficiency were xanthine stones. To test this, we looked for the presence of autofluorescent stones in animals carrying the *ok3234* null mutation in *xdh-1,* the *C. elegans* orthologue of xanthine dehydrogenase. In *C. elegans,* XDH-1 is expressed in the intestine, excretory cell, and neurons [21]. When we cultured *xdh-1* null mutant *C. elegans* on wild-type *E. coli* we indeed observed the formation of highly autofluorescent stones in 2% of animals (**Fig. 1B,C**). Thus, *xdh-1* was necessary for inhibiting the formation of autofluorescent stones. Consistent with their presence in diverse models of XDH-1-deficiency, we propose that the autofluorescent stones we observe during Moco- and XDH-1-deficiency in *C. elegans* are xanthine stones.

### *sulp-4* inhibited the formation of xanthine stones

XDH-1 functions at the end of the purine catabolism pathway to oxidize hypoxanthine to xanthine and xanthine to uric acid (**Fig. 1A**). Given the critical position of XDH-1 in purine metabolism, we were surprised that only 2% of *xdh-1* null mutant animals developed a xanthine stone. This result suggests the existence of parallel pathways for maintaining purine homeostasis. To identify additional regulators of purine metabolism, we performed an unbiased chemical mutagenesis screen for mutations that enhanced the penetrance of the xanthine stone phenotype. We mutagenized *xdh-1* mutant *C. elegans* with ethyl methanesulfonate (EMS) and cultured the newly mutagenized animals for 2 generations allowing newly induced mutations to become homozygous [22]. We then cloned single mutagenized F2 animals onto their own petri dish and screened for clones where we observed a high fraction of F3 progeny developing xanthine stones.

Here we describe five new EMS-induced recessive loss-of-function mutations that caused a high penetrance of xanthine stone formation in an *xdh-1* mutant background, *rae299, rae302, rae319, rae320* and *rae326* (**see Methods**). These mutant alleles were prioritized because they displayed strong enhancement of xanthine stone formation and formed a complementation group, indicating they affect a single gene (**see Methods**). To identify the causative genetic lesion in these new mutant strains, genomic DNA from all five strains was analyzed via whole genome sequencing. Our complementation studies suggested that the mutant strains should have novel mutations in a common gene. Only one gene, *sulp-4,* was uniquely mutated in all five strains, strongly suggesting these mutations were causative for the enhanced penetrance of xanthine stone formation in the *xdh-1* mutant animals (**Fig. 2A,B, Table S1**). Among the newly isolated *sulp-4* alleles we found 3 missense and 2 splice site mutations. Based on their recessive nature and molecular identities, we propose that these are loss-of-function alleles of *sulp-4*.

**Figure 2:**
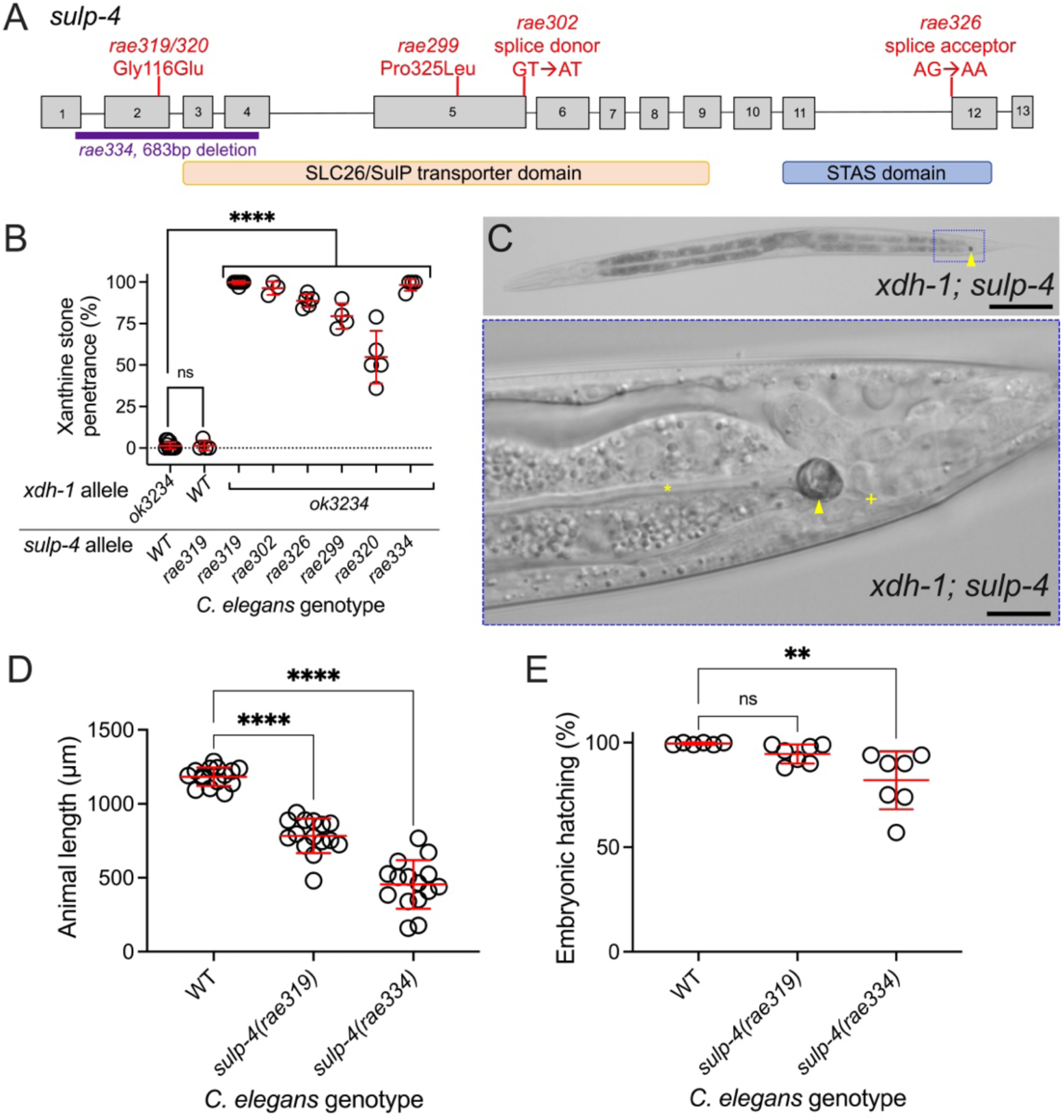
Loss-of-function mutations in *sulp-4* enhanced the formation of xanthine stones during *xdh-1* deficiency. A) *sulp-4* locus. Gray boxes are exons and lines are introns. The regions that encode the SLC26/SulP sulfate permease domain (orange) and STAS domain (blue) are displayed. Red lines display the location of new EMS-induced lesions that enhanced the formation of xanthine stones in an *xdh-1(ok3234)* mutant background. *rae334* (purple) is a deletion allele generated using CRISPR/Cas9 technology. B) Xanthine stone formation was assessed for *sulp-4(rae319)* single mutant and *xdh-1(ok3234)* mutant *C. elegans* with wild-type (WT) or mutant *sulp-4(rae319, rae302, rae326, rae299, rae320,* or *rae334).* ****, p<0.0001, ns, p>0.05, ordinary one-way ANOVA. Individual data points represent biological replicates. Mean and standard deviation are displayed. Complete information regarding sample size and individuals scored per biological replicate are found in **Table S2**. C) Differential interference contrast image of *xdh-1(ok3234); sulp-4(rae319) C. elegans* at the L4 stage of development. Blue box indicates the region magnified in the lower panel. Yellow arrowhead identifies the xanthine stone. Yellow asterisk identifies the lumen of the intestine. Yellow plus sign identifies the rectum. Scale bars are 100μm (top) and 10μm (bottom). D) Wild-type, *sulp-4(rae319),* and *sulp-4(rae334)* mutant *C. elegans* were synchronized at the first stage of larval development and cultured for 72 hours on wild-type *E. coli.* Animal length was determined. Individual datapoints are displayed as are the mean and standard deviation. The sample size is 15 individuals per genotype. ****, p<0.0001, ordinary one-way ANOVA. E) The hatching rate of newly laid wild-type, *sulp-4(rae319),* and *sulp-4(rae334)* mutant *C. elegans* embryos was determined. Individual data points represent biological replicates. Mean and standard deviation are displayed. Complete information regarding sample size and individuals scored per biological replicate are found in **Table S2**. **, p<0.01 or ns, p>0.05, ordinary one-way ANOVA.

To test the hypothesis that loss of *sulp-4* function causes enhanced xanthine stone accumulation in an *xdh-1* mutant, we used CRISPR/Cas9 to engineer a new *sulp-4* deletion allele, *rae334* [23, 24]. The *sulp-4(rae334)* allele is a 683 bp deletion that eliminates part of exon 1, all of exons 2 and 3, and part of exon 4 (**Fig. 2A**). Thus, we propose that *sulp-4(rae334)* encodes a null allele. The *sulp-4(rae334)* allele strongly enhanced the penetrance of xanthine stone formation in *xdh-1* mutant *C. elegans,* phenocopying the *sulp-4* alleles isolated in our EMS screen (**Fig. 2B**). These data demonstrate that the *sulp-4* lesions identified by whole genome sequencing cause xanthine stone accumulation in *xdh-1* mutant animals. Furthermore, these data show that *sulp-4* acts in parallel with *xdh-1* to inhibit the accumulation of xanthine stones.

The xanthine stones observed in *xdh-1; sulp-4* double mutant animals localized to the posterior of the *C. elegans* intestinal lumen, consistent with the localization of the stones observed in *xdh-1* mutant animals (**Fig. 2C**). In addition to observing xanthine stones at a higher frequency in *xdh-1; sulp-4* mutant animals, the xanthine stones were also much larger suggesting that *sulp-4* loss of function enhances both the penetrance and expressivity of the xanthine stone phenotype in *xdh-1* mutant animals (**Fig. S1**). This may reflect an increased quantity of xanthine in stones displayed by *xdh-1; sulp-4* mutant animals when compared to *xdh-1* single mutant animals.

We originally observed the formation of xanthine stones during conditions of Moco deficiency; *cth-2; moc-1* double mutant animals feeding on Moco-*E. coli.* We hypothesized that the *sulp-4* mutation would also enhance the formation of xanthine stones caused by Moco deficiency. To test this, we assayed xanthine stone formation in *sulp-4* single mutant animals cultured on wild-type or Moco-*E. coli*. 89% of *sulp-4* mutant animals developed xanthine stones during dietary Moco deficiency compared to 1% of *sulp-4* mutant animals fed a Moco replete diet (**Fig. S2A**). These data are consistent with our conclusion that *sulp-4* functions to limit the accumulation of xanthine stones caused by XDH-1 inactivation resulting from either Moco insufficiency or an *xdh-1* mutation.

### *sulp-4* promoted healthy larval and embryonic development

While culturing the *sulp-4(rae334)* mutant strain, we observed that mutant animals were sick and slow growing. Thus, we explored the role of *sulp-4* in development and embryonic viability. To test the impact of *sulp-4* loss of function on developmental rate, we synchronized wild-type, *sulp-4(rae319),* and *sulp-4(rae334)* animals at the first stage of larval development and assayed their growth after 72 hours. We found that *sulp-4(rae334)* animals displayed a severe developmental delay compared to the wild type (**Fig. 2D**). After 72 hours growth from L1, 100% of wild type animals reached the young adult stage (*n*=21), Only 5% of *sulp-4(rae334)* animals reached the young adult stage with 47.5% reaching L4 and the remaining 47.5% failing to reach L4 (*n*=24). Interestingly, *sulp-4(rae319)* animals showed a more subtle developmental delay (38% young adult, 58% L4, 4% pre-L4, *n*=21, **Fig. 2D**). Thus, we propose that *sulp-4(rae334)* is a null allele while *sulp-4(rae319)* represents a hypomorph. Similarly, we observed that *sulp-4(rae334)* caused 18% embryonic lethality while *sulp-4(rae319)* caused 5% embryonic lethality. No embryonic lethality was observed for wild-type *C. elegans* (**Fig. 2E**). Thus, we conclude that *rae334* and *rae319* represent an allelic series for *sulp-4* and that *sulp-4* is necessary for promoting embryonic and larval development in *C. elegans*.

### *pnp-1* was necessary for the formation of xanthine stones in *xdh-1; sulp-4* mutants animals

To further test the model that the autofluorescent stones observed in *xdh-1; sulp-4* mutant animals were composed of xanthine, we performed genetic epistasis with a null mutation in purine nucleoside phosphorylase (*pnp-1*), a gene necessary for the formation of hypoxanthine and xanthine (**Fig. 1A**) [25]. PNP-1 is expressed and acts in the *C. elegans* intestine in addition to expression in some head neurons [25]. As previously observed, *xdh-1; sulp-4* double mutant animals displayed 98% autofluorescent stone formation while *pnp-1 xdh-1; sulp-4* triple mutant *C. elegans* displayed 3% formation of autofluorescent stones (**Fig. 3A**). Thus, *pnp-1* was necessary for the formation of the autofluorescent stones observed in *xdh-1; sulp-4* double mutant animals. Given that *pnp-1* plays a conserved role in the formation of hypoxanthine and xanthine, these results support our model that the autofluorescent stones we observe are likely to be predominantly composed of xanthine. Although, we cannot exclude the possibility that other metabolites, such as hypoxanthine, are also present in the autofluorescent stones.

**Figure 3:**
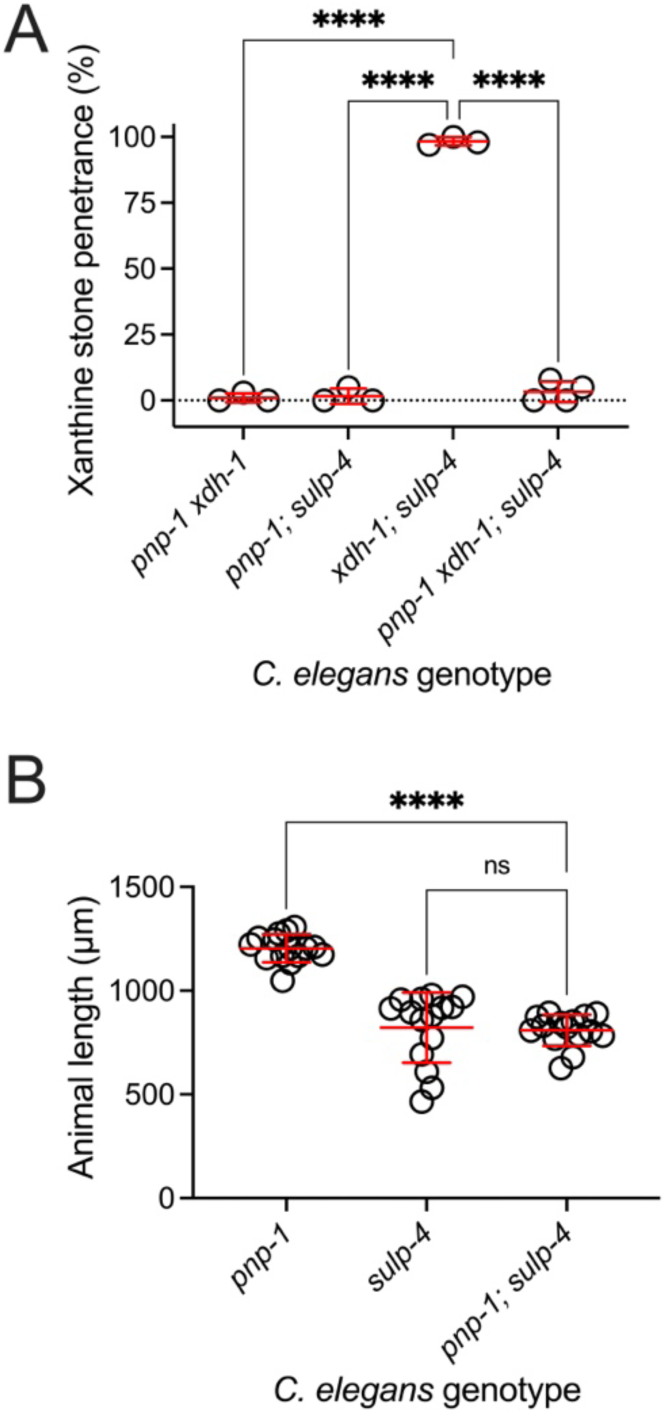
*pnp-1* was required for xanthine stone formation displayed by *xdh-1; sulp-4* mutants but not the larval delay caused by *sulp-4* loss of function. A) Double and triple mutant *C. elegans* were assessed for xanthine stone formation when cultured on wild-type *E. coli*. ****, p<0.0001, ordinary one-way ANOVA. Individual data points represent biological replicates. Mean and standard deviation are displayed. Complete information regarding sample size and individuals scored per biological replicate are found in **Table S2**. B) *pnp-1(jy121), sulp-4(rae319),* and *pnp-1(jy121); sulp-4(rae319)* mutant *C. elegans* were synchronized at the first stage of larval development and cultured for 72 hours on wild-type *E. coli.* Animal length was determined. Individual datapoints are displayed as are the mean and standard deviation. The sample size is 15 individuals per genotype. ****, p<0.0001 or ns, p>0.05, ordinary one-way ANOVA.

To determine if *pnp-1* acts in a genetic pathway with *sulp-4,* we tested the impact of *pnp-1* loss of function on the developmental delay displayed by *sulp-4* mutant animals. *pnp-1; sulp-4* double mutant larvae developed at a rate similar to *sulp-4* single mutant animals (**Fig 3B**). After 72 hours growth from L1, 81% of *pnp-1; sulp-4* double mutant animals reached the young adult stage while the remaining 19% reached L4 (*n*=31). Similarly, 83% of *sulp-4* single mutant animals reached young adult, 14.5% reached L4 and 2.5% failed to reach L4 (*n*=42). Importantly, *pnp-1* mutant animals displayed healthy larval development (100% young adults 72 hours post L1, n=39, **Fig. 3B**). Thus, *pnp-1* was not required for the developmental delay displayed by *sulp-4* mutant *C. elegans.* We propose a genetic pathway where *pnp-1* promotes the formation of xanthine stones epistatic to the function of *xdh-1* and in parallel to the activity of *sulp-4*.

### *sulp-*4/SLC26 encodes a sulfate permease that acted in the excretory cell to promote xanthine homeostasis

*sulp-4* encodes a transmembrane transporter with homology to the SLC26 family of anion transporters in mammals [13, 26]. The *C. elegans* genome encodes eight members of the SLC26 transporter family, named SULP-1 through SULP-8 [13]. We wondered if other members of the SLC26 family of transporters also played a role in limiting the formation of xanthine stones. To test this, we cultured strains with deletions in *sulp-1, sulp-2, sulp-4, sulp-5, sulp-7,* and *sulp-8* on Moco-*E. coli* and assayed the formation of xanthine stones. *sulp-3* and *sulp-6* mutant strains were not analyzed as they are inviable. Only the strain carrying the *sulp-4* mutation displayed a high penetrance of xanthine stones (88%, **Fig. S2B**). Thus, the enhancement of xanthine stone formation is specific to loss of *sulp-4* and not a general feature of *sulp* inactivation.

SULP-4 is expressed in the apical membrane of the *C. elegans* excretory cell, a single cell that plays roles in ionic regulation and waste elimination analogous to the mammalian renal system [13, 14, 27]. Studies of SULP-4 expressed in *Xenopus* oocytes demonstrate that SULP-4 is sufficient to promote the transport of sulfate and, to a lesser extent, chloride [13]. To determine the site of action of *sulp-4* with respect to the xanthine stone formation phenotype, we generated transgenic *xdh-1; sulp-4* double mutant *C. elegans* expressing a *Psulp-4::SULP-4::GFP* translational fusion transgene (plasmid was a gift from Dr. Keith Nerhke) [13]. Consistent with previous reports, we exclusively saw expression of the *Psulp-4::SULP-4::GFP* translational fusion in the excretory cell (**Fig. 4A,B**). To test the functionality of the *Psulp-4::SULP-4::GFP* transgene, we performed rescue experiments with *xdh-1; sulp-4* double mutant animals expressing *Psulp-4::SULP-4::GFP* and assaying the formation of xanthine stones. *xdh-1; sulp-4* double mutant animals expressing *Psulp-4::SULP-4::GFP* did not develop xanthine stones, demonstrating functional transgenic rescue. This rescue was observed in three independently derived transgenic strains (**Fig. 4C**). Thus, the *Psulp-4::SULP-4::GFP* transgenic fusion protein was functional, suggesting that its expression pattern faithfully represents endogenous SULP-4 localization. We conclude that SULP-4 acts in the excretory cell to negatively regulate the formation of xanthine stones. Our observation that *xdh-1; sulp-4* double mutant animals develop xanthine stones in the intestinal lumen suggests that *sulp-4* is functioning cell non-autonomously to limit the formation of xanthine stones.

**Figure 4:**
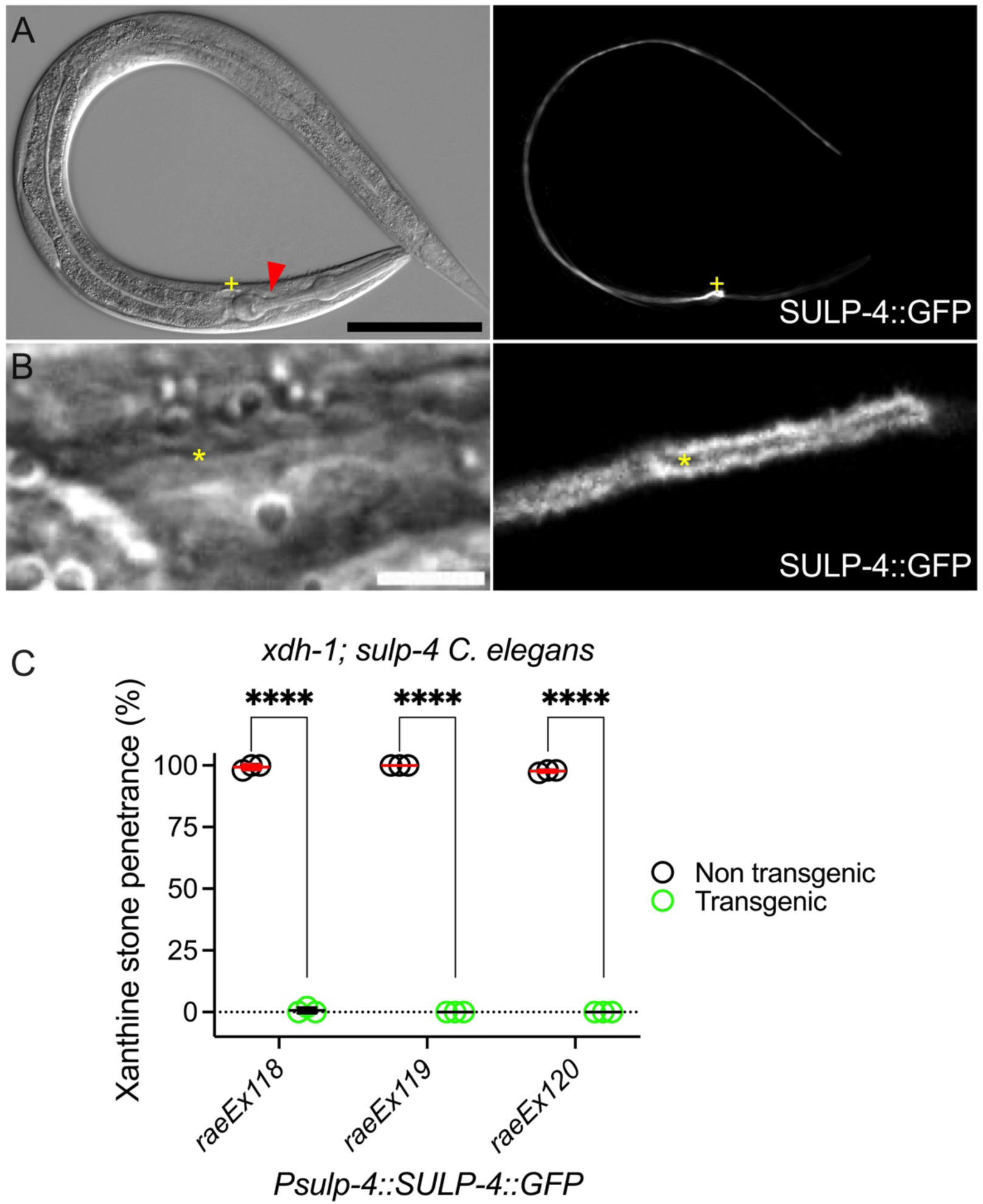
*sulp-4* acted cell non-autonomously in the excretory cell to limit xanthine stone formation. A,B) Differential interference contrast (left) and fluorescence imaging (right) of *xdh-1(ok3234); sulp-4(rae319) C. elegans* expressing the *Psulp-4::SULP-4::GFP* transgene (SULP-4::GFP). Yellow plus sign identifies the cell body of the excretory cell. Red arrow indicates the region magnified in panel B. Yellow asterisk identifies the lumen of the excretory cell. Scale bars are 100μm (A, top) and 5μm (B, bottom). C) Transgenic *xdh-1(ok3234); sulp-4(rae319) C. elegans* expressing the *Psulp-4::SULP-4::GFP* transgene (green) and their non-transgenic siblings (black) were assessed for the formation of xanthine stones. Individual data points represent biological replicates. Mean and standard deviation are displayed. ****, p<0.0001, multiple unpaired t tests. Complete information regarding sample size and individuals scored per biological replicate are found in **Table S2**.

To further test the role of the excretory cell in preventing the formation of xanthine stones, we used an *exc-5* mutation that causes defects in excretory cell development and morphology [28]. We reasoned a malformed excretory cell may not function efficiently and thus phenocopy *sulp-4* loss of function with respect to xanthine stone formation. Indeed, *exc-5; xdh-1* double mutant animals displayed enhanced formation of xanthine stones (**Fig. S3A**). Although, the xanthine stone penetrance of the *exc-5; xdh-1* (17%) double mutant strain was modest compared to *xdh-1; sulp-4*. Importantly, *sulp-4* loss of function did not cause cysts in the excretory cell tubules, a severe defect in excretory cell morphology caused by *exc-5* loss of function (**Fig. S3B-D**). Although, we cannot exclude a subtle defect in excretory cell tubule extension in *sulp-4* mutant animals. We conclude that the enhancement of xanthine stone formation caused by inactivating mutations in *sulp-4* result from loss of SULP-4 anion exchange function and not broader defects in excretory cell biology.

Furthermore, ion homeostasis plays a key role in the defecation cycle which promotes waste elimination from the *C. elegans* intestine [29]. We wondered if the enhanced formation of xanthine stones observed in *xdh-1; sulp-4* animals might result from failures in defecation, leading to xanthine accumulation in the intestinal lumen and xanthine stone formation. To test this model, we used the *aex-5(sa23)* mutation which causes defects in anterior body-wall muscle contraction (aBoc) and waste expulsion (Exp), critical aspects of the defecation cycle [30]. Interestingly, *aex-5; xdh-1* double mutant animals did not display an increased penetrance of xanthine stone formation (**Fig. S3A**). These results suggest that defects in defecation alone are not sufficient to promote xanthine stone accumulation.

### *cth-2* and *cdo-1* were necessary for *sulp-4* mutant phenotypes

*sulp-4* inactivation caused xanthine stone formation during dietary Moco deficiency (**Fig. S2A, Fig. S4**), demonstrating that the enhancement of xanthine stones caused by *sulp-4* loss of function occurs even when endogenous Moco biosynthesis is functional. To test the impact of a *sulp-4* mutation on xanthine stone formation during complete Moco deficiency, we engineered *sulp-4; cdo-1 moc-1* triple mutant *C. elegans* that cannot synthesize their own Moco (caused by *moc-1* mutation) and are viable during Moco deficiency (caused by *cdo-1* suppressor mutation) (**Fig. 1A**). When cultured on Moco-*E. coli, sulp-4; cdo-1 moc-1* triple mutants are completely Moco deficient yet only displayed 9% penetrance of xanthine stones (**Fig. S4**). Similarly, *sulp-4; cdo-1* double mutant animals cultured on Moco-*E. coli* displayed 0% penetrance of xanthine stones, dramatically reduced compared to the 86% penetrance displayed by *sulp-4* mutant animals cultured on Moco-*E. coli* (**Fig. S4**). These results were surprising and suggest that *cdo-1* is necessary for the formation of xanthine stones caused by a *sulp-4* mutation and Moco deficiency.

To further test the impact of *cdo-1* on the formation of xanthine stones, we engineered *xdh-1; sulp-4; cdo-1* triple mutant *C. elegans*. Surprisingly, *xdh-1; sulp-4; cdo-1* triple mutant animals displayed a 5% xanthine stone penetrance, dramatically reduced when compared to the 98% penetrance displayed by *xdh-1; sulp-4* double mutant animals (**Fig. 5A**). Thus, *cdo-1* was necessary for the formation of xanthine stones displayed by *xdh-1; sulp-4* double mutant animals. Interestingly, we still observe a low penetrance of xanthine stones in *xdh-1; sulp-4; cdo-1* triple mutant *C. elegans* suggesting that *cdo-1* activity is not absolutely required for the formation of xanthine stones but only required for the xanthine stone enhancement caused by *sulp-4* loss of function.

**Figure 5:**
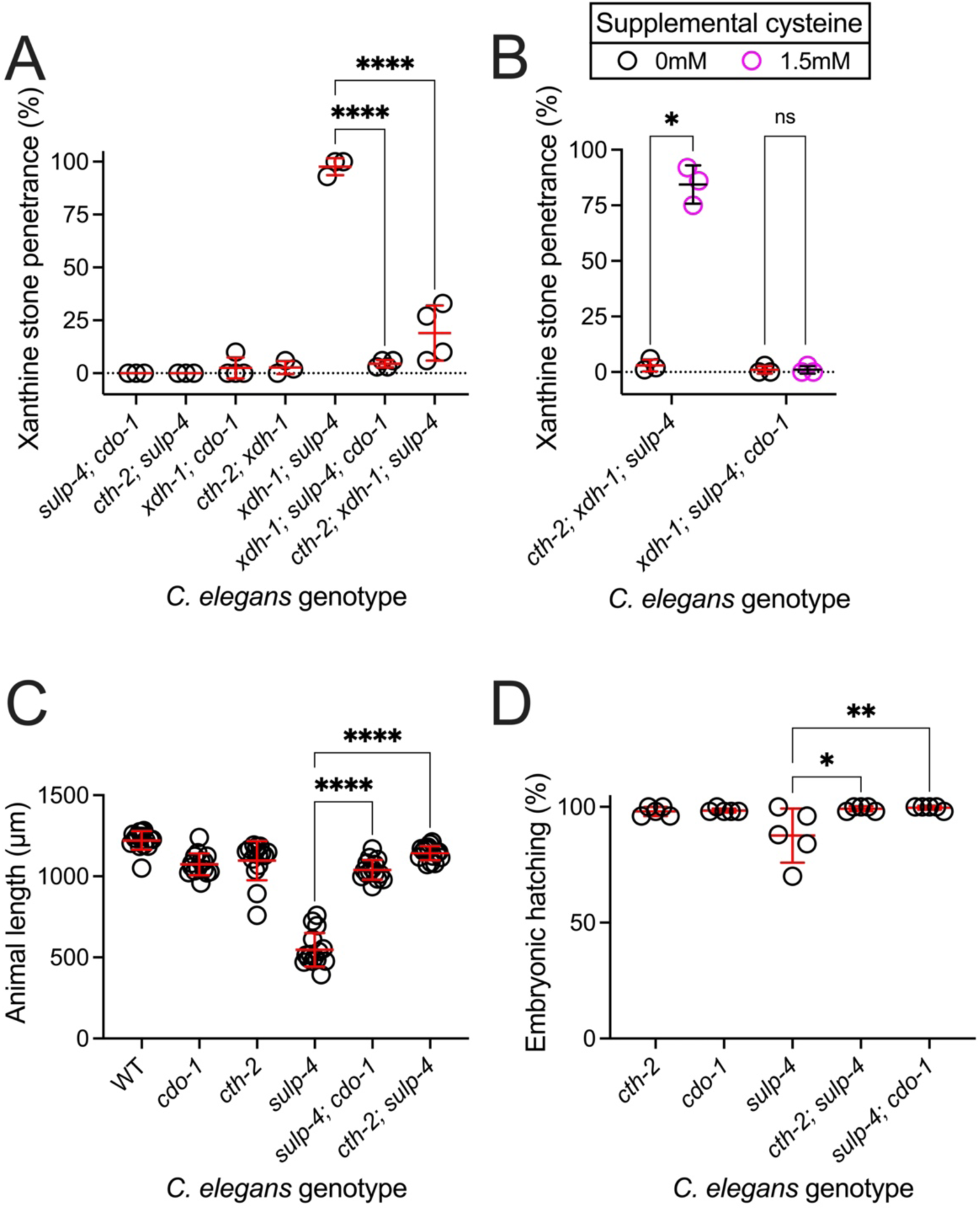
*cth-2* and *cdo-1* were required for phenotypes caused by *sulp-4* loss of function. A) Double and triple mutant *C. elegans* were assessed for xanthine stone formation when cultured on wild-type *E. coli* over the first 3 days of adulthood. Individual data points represent biological replicates. Mean and standard deviation are displayed. ****, p<0.0001, ordinary one-way ANOVA. Complete information regarding sample size and individuals scored per biological replicate are found in **Table S2**. B) Triple mutant *C. elegans* exposed to 0 or 1.5mM supplemental cysteine were assessed for xanthine stone formation when cultured on wild-type *E. coli* over the first 2 days of adulthood. Mean and standard deviation are displayed. *, p<0.05 and ns, p>0.05, multiple unpaired t tests. Complete information regarding sample size and individuals scored per biological replicate are found in **Table S2**. C) Wild-type and mutant *C. elegans* were synchronized at the first stage of larval development and cultured for 72 hours on wild-type *E. coli.* Animal length was determined. Individual datapoints are displayed as are the mean and standard deviation. The sample size is 15 individuals per genotype. ****, p<0.0001, ordinary one-way ANOVA. D) The hatching rate of newly laid single and double mutant *C. elegans* embryos was determined. Individual data points represent biological replicates. Mean and standard deviation are displayed. **, p<0.01 or *, p<0.05, ordinary one-way ANOVA. Complete information regarding sample size and individuals scored per biological replicate are found in **Table S2**. The *sulp-4(rae334)* allele was used to generate the data in Fig. 5.

*cdo-1* encodes the *C. elegans* cysteine dioxygenase, a critical enzyme in the in sulfur amino acid catabolism pathway that breaks down excess cysteine and methionine (**Fig. 1A**) [8, 31, 32]. To determine if the impact of *cdo-1* on the enhanced xanthine stone formation caused by *sulp-4* inactivation was a result of impaired sulfur amino acid catabolism, we used a *cth-2* mutation which eliminates the activity of *C. elegans* cystathionase (**Fig. 1A**). Consistent with our results with *cdo-1* loss of function, we found that *cth-2; xdh-1; sulp-4* triple mutant animals also displayed a low 19% penetrance of xanthine stones (**Fig. 5A**). Taken together, these genetic data suggest that sulfur amino acid catabolism is required for the enhancement of xanthine stone accumulation caused by loss of *sulp-4* function.

Given that *sulp-4* encodes a sulfate permease, we hypothesized that sulfate accumulation during *sulp-4* loss of function was promoting the formation of xanthine stones. Furthermore, we hypothesized that this sulfate was produced endogenously via CTH-2 and CDO-1. To further test this model, we measured sulfur content in large cultures of wild-type and *sulp-4* mutant *C. elegans* using inductively coupled plasma mass spectrometry (ICP-MS). We did not observe any difference in sulfur content between wild-type and *sulp-4* mutant animals, demonstrating that *sulp-4* mutant *C. elegans* do not accumulate sulfur (**Fig. S5B**). Similarly, we saw no difference in K, Fe, Zn, Mn, Cu or Mo content between wild type and *sulp-4* mutant animals (**Fig. S5**). A key limitation of this method is that ICP-MS does not distinguish between sulfur found in different metabolites such as sulfate, sulfite, methionine, cysteine, etc. Thus, it is plausible that *sulp-4* mutant animals accumulate sulfate with a corresponding decrease in another sulfur-containing metabolite. Alternatively, sulfate sulfur may only account for a small fraction of the animal sulfur economy. Thus, physiologically significant changes in sulfate may be undetectable given the high background of sulfur found in more common metabolites like cysteine or methionine.

To further test the relationship between sulfur amino acid catabolism and xanthine stone formation, we supplemented *xdh-1; sulp-4; cdo-1* or *cth-2; xdh-1; sulp-4* triple mutant *C. elegans* with the sulfur-containing amino acid cysteine and assessed the impact on xanthine stone formation. Cysteine supplementation is informative as it is the product of CTH-2 and the substrate for CDO-1 (**Fig. 1A**). We found that cysteine promoted xanthine stone formation in *cth-2; xdh-1; sulp-4* animals, bypassing the requirement for *cth-2.* Alternatively, cysteine did not promote xanthine stones in *xdh-1; sulp-4; cdo-1* mutant animals, demonstrating that *cdo-1* was necessary for cysteine to promote xanthine stone formation (**Fig. 5B**). This result supports our model that endogenous sulfur amino acid catabolism promotes xanthine stone formation. More specifically, these data demonstrate that a cysteine-derived metabolite downstream of CDO-1 is acting to promote xanthine stones. This is consistent with our model that sulfate accumulation during *sulp-4* loss of function promotes xanthine stone accumulation.

To test whether mutations in *cth-2* or *cdo-1* would suppress the defects in larval and embryonic development displayed by *sulp-4* mutant animals, we assayed larval and embryonic development in *cth-2; sulp-4* and *sulp-4; cdo-1* double mutant animals and compared to *sulp-4* single mutant animals. Consistent with their suppression of xanthine stone formation, *cth-2* or *cdo-1* mutations suppressed the developmental delay and embryonic lethality caused by a *sulp-4* mutation (**Fig. 5C,D**). After 72 hours growth from L1, 0% of *sulp-4* mutant animals reached the young adult stage, 53% reached L4, and 47% failed to reach L4 (*n*=32). In contrast, 100% of *cth-2; sulp-4* or *sulp-4; cdo-1* double mutant animals reached the young adult stage (*n*=32) Thus, we conclude that *cth-2* and *cdo-1* are broadly required for phenotypes caused by *sulp-4* loss of function.

### Loss of *osm-8* promoted xanthine stone accumulation in *xdh-1* mutant animals, linking osmotic homeostasis and xanthine stone formation

What is the mechanism by which sulfates, derived from CTH-2 and CDO-1, might promote xanthine stone formation when *sulp-4* is inactivated? Given that *sulp-4* encodes a sulfate permease that functions in the *C. elegans* excretory cell, we hypothesized that *sulp-4* mutant animals may be experiencing osmotic imbalance driven by excess sulfate. Supporting this model, *sulp-4* mutant animals accumulated 25% more sodium than their wild-type counterparts as measured by ICP-MS (**Fig. 6A**). These data demonstrate that *sulp-4* mutant animals are experiencing osmotic imbalance.

**Figure 6:**
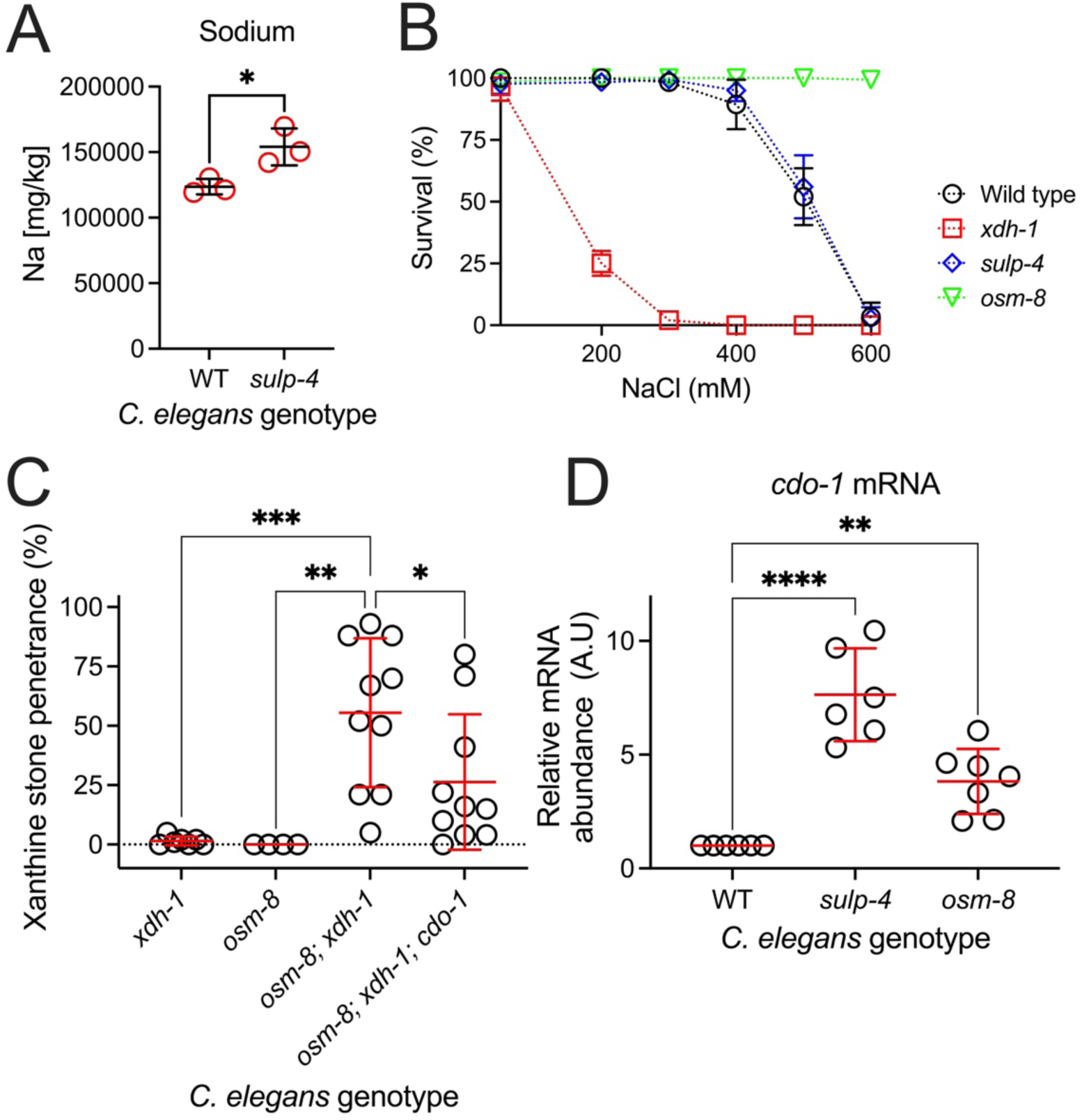
Loss of *osm-8,* which activates the osmotic stress response, enhanced xanthine stone accumulation in *xdh-1* mutant *C. elegans*. A) Sodium content was determined for synchronized wild-type and *sulp-4(rae319)* young adult *C. elegans* using ICP-MS. Individual data points represent biological replicates. Mean and standard deviation are displayed. *, p<0.05, t test. B) Survival of wild-type, *sulp-4(rae319), xdh-1(ok3234),* or *osm-8(n1518)* young adult *C. elegans* was determined after 24-hour exposure to NGM media containing various concentrations of NaCl. Mean and standard deviation of 3 biological replicates are displayed. Complete information regarding sample size and individuals scored per biological replicate are found in **Table S2**. C) Mutant *C. elegans* were assessed for xanthine stone formation when cultured on wild-type *E. coli*. Individual data points represent biological replicates. Mean and standard deviation are displayed. ***, p<0.001, **, p<0.01, *, p<0.05, ordinary one-way ANOVA. Complete information regarding sample size and individuals scored per biological replicate are found in **Table S2**. D) Relative mRNA levels of *cdo-1* are displayed for RNA isolated from wild-type, *sulp-4(rae319)*, and *osm-8(n1518)* young adult *C. elegans.* Relative mRNA abundance was determined via the delta-delta C_T_ method. All transcripts are normalized to *act-1.* Relative mRNA abundance for each transcript was set to one in the wild type. ****, p<0.0001, **, p<0.01, ordinary one-way ANOVA.

To test the model that osmotic imbalance promotes the formation of xanthine stones, we challenged *xdh-1* mutant animals with supplemental NaCl. Interestingly, *xdh-1* mutant animals were extremely sensitive to high environmental NaCl when compared to wild type, a phenotype also displayed by rats lacking *xdh-1* (**Fig. 6B**) [33]. These data indicate a conserved role for *xdh-1* in promoting hyperosmotic stress tolerance. However, the sensitivity of *xdh-1* mutant *C. elegans* to NaCl limited our ability to directly test whether hyperosmotic stress promotes xanthine stone formation in an *xdh-1* mutant background.

To genetically test whether osmotic imbalance may promote xanthine stones, we used a loss-of-function mutation in *osm-8* that constitutively activates the hyperosmotic stress response in the absence of environmental osmotic stress [15, 34]. *osm-8* encodes a mucin-like protein and is expressed in the *C. elegans* hypodermis [15]. Interestingly, *osm-8; xdh-1* double mutant *C. elegans* developed xanthine stones like *xdh-1; sulp-4* double mutant animals (**Fig. 6C)**. Although, the xanthine stones observed in *osm-8; xdh-1* double mutant animals were not as large as those observed in *xdh-1; sulp-4* animals (**Fig. S1).** Furthermore, like *xdh-1; sulp-4* double mutant *C. elegans*, *cdo-1* was necessary for the formation of xanthine stones in *osm-8; xdh-1* double mutant animals (**Fig. 6C)**. However, it should be noted that *cdo-1* loss of function incompletely suppressed the formation of xanthine stones in *osm-8; xdh-1* mutant animals, suggesting additional pathways downstream of *osm-8* that govern xanthine stone formation. These data suggest that altered osmotic homeostasis, driven by *osm-8* loss of function, is sufficient to cause xanthine stone accumulation when XDH-1 is inactive.

We wondered if *sulp-4* and *osm-8* mutant animals display additional overlapping phenotypes. *osm-8* mutant *C. elegans* are resistant to high NaCl stress and display increased transcription of the hypertonic stress response which includes the key regulators of osmolyte accumulation, *gpdh-1* and *hmit-1.1* [15, 34]. *gpdh-1* encodes glycerol 3-phosphate dehydrogenase and is required to produce glycerol, a critical osmolyte [35, 36]. *hmit-1.1* encodes an H^+^/*myo-*inositol transporter and functions to import *myo-*inositol, another important osmolyte [37]. Distinct from *osm-8* mutant animals, *sulp-4* mutant *C. elegans* were not resistant to high NaCl (**Fig. 6B)**. Furthermore, *sulp-4* mutant animals did not significantly accumulate *hmit-1.1* mRNA and modestly accumulated *gpdh-1* mRNA when compared to the wild type **(Fig. S6A,B**). These data suggest that *sulp-4* inactivation does not robustly activate transcription of all canonical hypertonic stress response genes. Interestingly, both *sulp-4* and *osm-8* mutant animals accumulated *cdo-1* mRNA when compared to the wild type (**Fig. 6D**), consistent with previous studies in *C. elegans* and mammals that demonstrate *cdo-1* induction by osmotic imbalance (high NaCl exposure or *osm-8* inactivation) [34, 38]. Taken together, these data highlight overlapping but distinct phenotypic profiles for *sulp-4* and *osm-8* mutant *C. elegans* and elucidate a positive feedback loop connecting *sulp-4,* osmotic imbalance, and *cdo-1* activation.

In addition to its role in cysteine catabolism, CDO-1 is also essential for synthesis of the osmolyte taurine [32, 38]. Given that *cdo-1* was necessary for phenotypes caused by *sulp-4* inactivation, we wondered whether inactivating mutations in other genes involved in osmolyte accumulation may also suppress *sulp-4* mutant phenotypes. To test this, we engineered *gpdh-1; sulp-4* and *hmit-1.1; sulp-4* double mutant strains of *C. elegans* and assayed for the formation of xanthine stones when animals were cultured on a Moco-diet. Unlike *cdo-1,* neither *gpdh-1* nor *hmit-1.1* were necessary for the formation of xanthine stones caused by a *sulp-4* mutation (**Fig. S6C**). These data suggest that the suppression of *sulp-4* mutant phenotypes by *cdo-1* loss-of-function is specific and not a general feature of inactivating mutations in genes involved in osmolyte accumulation.

We propose the model that in healthy wild-type *C. elegans,* cysteine and methionine are being broken down by the sulfur amino acid catabolism pathway (CTH-2/CDO-1), maintaining sulfur homeostasis. This results in the production of sulfate which is exported into the environment via the action of SULP-4 in the excretory cell. However, when *sulp-4* is inactive, sulfate cannot be safely excreted and accumulates. This issue is exacerbated by an induction of *cdo-1* mRNA during *sulp-4* loss of function. This creates a maladaptive positive feedback loop that we anticipate would further increase sulfate production. We speculate that sulfate accumulation then promotes osmotic imbalance which causes embryonic lethality, impaired larval development, and an increased propensity to form xanthine stones when *xdh-1* is inactive (**Fig. 7**).

**Figure 7:**
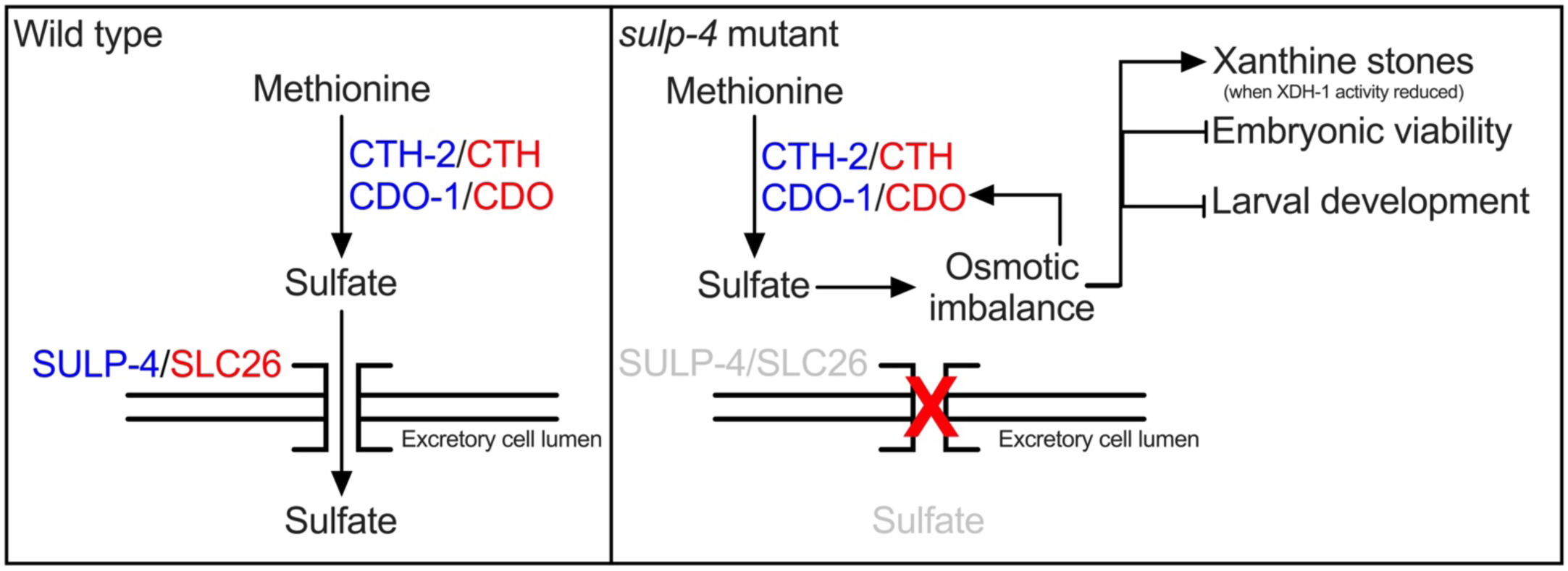
SULP-4 promotes sulfate homeostasis, maintaining osmotic balance in *C. elegans*. In wild-type *C. elegans*, sulfur amino acid catabolism gives rise to sulfate which is maintained at homeostatic levels by SULP-4-mediated exchange with the environment via the excretory cell. During *sulp-4* loss of function, CTH-2/CDO-1-derived sulfates accumulate causing osmotic imbalance. This osmotic imbalance promotes a maladaptive positive feedback loop promoting additional *cdo-1* mRNA accumulation. This cascade culminates in embryonic lethality, altered larval development, and a propensity to form xanthine stones when XDH-1 activity is compromised.

## Discussion

### Modeling xanthinuria in C. elegans

Human xanthinuria was originally described in 1954, and presents with high urinary xanthine, low uric acid in serum and urine, the formation of xanthine stones, and, in some cases, renal failure [5, 39]. Still, there are no curative treatments for xanthinuria or the formation of xanthine stones. The current recommendation for patients is a high fluid intake and low purine diet [7]. Thus, understanding the cellular mechanisms that regulate the pathology associated with xanthinuria is an important goal.

Animal models, such as *C. elegans,* are powerful tools for exploring the pathology of rare inborn errors of metabolism including NGLY1 deficiency, Moco deficiency, Friedrich’s ataxia, and many others [8, 40, 41]. Here we used *C. elegans* to model human type I and type II xanthinuria. We employed genetic strategies to inhibit XDH-1 activity by mutating the *xdh-1* gene (type I) or limiting animal Moco (type II) [4, 5]. Both manipulations recapitulated a critical feature of human XDH deficiency, the formation of insoluble xanthine stones.

Interestingly, xanthine stones are highly autofluorescent and visible with a standard fluorescence microscope, a phenotype that has been previously characterized in the model plant *Arabidopsis thaliana* [18]. Given the transparent nature of *C. elegans,* this phenotype empowers genetic analyses of xanthine stone accumulation and, by proxy, purine biology. Here we used the power of *C. elegans* genetics in combination with this simple phenotype to identify and characterize regulators of purine homeostasis.

### Defining genetic regulators of xanthine stone formation

We sought to define genetic regulators of the formation of xanthine stones. Given the established purine catabolism pathway (**Fig. 1A)**, we used a hypothesis-driven approach to define genes that regulate the formation of xanthine stones. Purine nucleoside phosphorylase (PNP-1/PNP) was a lead candidate given its biochemical requirement for the formation of xanthine. Indeed, we demonstrated that *pnp-1* was necessary for the formation of xanthine stones in our *C. elegans* mutant animals: *pnp-1* loss of function suppressed the formation of xanthine stones in an *xdh-1; sulp-4* double mutant background. These results suggest that inhibiting the activity of PNP may be a therapeutic strategy for limiting the accumulation of xanthine and xanthine stones in patients suffering from xanthinuria. However, this treatment strategy may be fraught, given the consequences of PNP inactivation. Human patients with PNP deficiency display impaired T-cell immunity [3]. In fact, a potent PNP inhibitor has been developed, and induces apoptosis of B- and T-lymphocytes [42, 43]. Thus, the potential benefits of PNP inhibition in the treatment of human xanthinuria patients would need to be evaluated and weighed against the negative impacts on the immune system.

To identify new and unexpected regulators of purine homeostasis, we employed an unbiased genetic approach. In a forward genetic screen, we identified *sulp-4* as a potent modifier of xanthine stone formation. Loss-of-function mutations in *sulp-4* dramatically enhanced the penetrance and expressivity of xanthine stone formation in our *C. elegans* models of xanthinuria. Interestingly, we also found that *sulp-4* was necessary for promoting normal larval and embryonic development. These genetic data demonstrate that *sulp-4* promotes healthy development and acts in a parallel pathway to *xdh-1* to limit the accumulation of xanthine stones.

*sulp-4* was previously demonstrated to encode a sulfate permease that localizes to the apical membrane of the excretory cell, which we also observed [13]. Importantly, we demonstrated that *Psulp-4::SULP-4::GFP* rescued the formation of xanthine stones displayed by an *xdh-1; sulp-4* double mutant animal. This functional rescue is strong evidence that *sulp-4* is acting in the excretory cell to limit the formation of xanthine stones. Interestingly, xanthine stones accumulate in the lumen of the *C. elegans* intestine while *sulp-4* acts in the excretory cell. Thus, we conclude that *sulp-4* acts cell-nonautonomously to limit xanthine stone accumulation. These data establish a new intersection between SULP-4, the excretory cell, and purine homeostasis.

### Linking osmotic homeostasis and xanthine stone formation

We next sought to understand the nature of the genetic interaction between *sulp-4* and *xdh-1* with respect to xanthine stone formation. Given that *sulp-4* encodes a sulfate transporter that acts cell-nonautonomously in the excretory cell, we hypothesized that *sulp-4* mutations may cause sulfate retention and disturb osmotic balance in *C. elegans* systemically [13]. Indeed, we found that *sulp-4* mutant *C. elegans* accumulated sodium, demonstrating osmotic imbalance [44, 45]. Importantly, we do not know the site of excess sodium accumulation (cellular vs. extracellular) in *sulp-4* mutant *C. elegans.* We propose the model that osmotic imbalance in *sulp-4* mutant animals is driven by a failure to excrete endogenously produced sulfates. This model is supported by our genetic demonstration that *cth-2* or *cdo-1,* genes that encode enzymes necessary for sulfate production, were necessary for xanthine stone formation in *xdh-1; sulp-4* mutant animals. We speculate that osmotic imbalance driven by sulfate retention may promote water reabsorption causing dehydration in the intestinal lumen, increasing xanthine concentration and stone formation. This model is reinforced by our observation that activation of the hyperosmotic stress response through a distinct genetic perturbation, *osm-8* loss of function, also promoted the formation of xanthine stones in an *xdh-1* mutant background [15]. In the case of *osm-8* inactivation, we speculate that glycerol accumulation would drive water reabsorption and dehydration in the intestinal lumen. Again, this dehydration would increase xanthine concentration, promoting stone formation.

### Intersecthion between sulfur amino acid metabolism and Moco-dependent metabolism

Previous studies in *C. elegans* identified *cth-2* and *cdo-1* loss of function mutations as suppressors of the lethality associated with Moco deficiency and deficiency of the Moco-requiring enzyme sulfite oxidase [8]. Loss of sulfite oxidase is lethal in *C. elegans* and humans due to the accumulation of its reactive and toxic substrate, sulfite (**Fig. 1A**) [8, 46]. *cth-2* and *cdo-1* inactivation limit the accumulation of sulfites, suppressing the lethality caused by Moco or sulfite oxidase deficiency. Is it a coincidence that loss of *cth-2* or *cdo-1* suppress phenotypes associated with two distinct Moco-requiring enzymes; i) lethality displayed by *suox-1* null mutant animals and ii) xanthine stone formation of *xdh-1; sulp-4* double mutant animals? Given that the substrates and products of these two Moco-dependent enzymes are distinct, it is peculiar that loss-of-function phenotypes of both are modulated by these common genetic factors. Future studies are required to tease apart the potential molecular intersections between CTH-2/CDO-1 and these Moco-dependent pathways.

An additional layer of complexity is added when considering the regulation of *cdo-1.* Here, we reinforce previous observations that *cdo-1* is activated in response to hyperosmotic stress [34, 38]. *sulp-4* and *osm-8* mutant *C. elegans* both accumulated *cdo-1* mRNA. Yet, CDO-1/CDO levels and activity are also modified by dietary sulfur amino acid content. This regulation includes both transcriptional and post-translational control of cysteine dioxygenase [31, 47–51]. Given that SULP-4 mediates sulfate transport and sulfate is a metabolic product of sulfur amino acid catabolism governed by CTH-2/CDO-1/SUOX-1, sulfate seems like a potential metabolic intersection between these seemingly disparate Moco-dependent pathways. Whether there is any overlap between the mechanisms of CDO-1/CDO regulation downstream of hyperosmotic stress and high sulfur amino acid content remains to be studied.

### SULP-4 homologs are implicated in calcium oxalate stone formation

Our genetic studies demonstrate that SULP-4 plays a role in limiting the formation of xanthine stones in *C. elegans* models of xanthinuria. SULP-4 is homologous to human SLC26 proteins which have been previously linked to metabolic stone formation. Understanding the connection between SLC26 proteins, like SULP-4, and the formation of metabolic stones is important as 1 out of every 10 individuals in the United States will develop a kidney stone over the course of their lifetimes [52]. For instance, mouse SLC26A6 limits urolithiasis, paralleling our results with *C. elegans* SULP-4. Specifically, SLC26A6 null mutant mice develop a high incidence of calcium oxalate stones in the bladder [53]. Similarly, a dominant negative mutation in SLC26A6 and loss of function mutations in SLC26A1 are proposed to cause calcium oxalate nephrolithiasis in humans [54, 55]. In contrast, the *Drosophila melanogaster* homolog of human SLC26A5/6, dPrestin, promotes the formation of calcium oxalate stones in the Malpighian tubules in a dietarily-induced model of calcium oxalate nephrolithiasis. Under a diet of high oxalate, *Drosophila* develop calcium oxalate stones whose formation is dampened by RNAi knockdown of dPrestin [56]. With respect to calcium oxalate stones, the mechanism of stone formation is believed to relate directly to the role of the SLC26 family members in oxalate transport. Loss of SLC26 proteins results in higher or lower concentrations of oxalate in a given space, altering the likelihood of stone formation. We propose that xanthine is the critical component of the stones that form in *xdh-1; sulp-4* mutant animals, not oxalate. In heterologous transport assays, oxalate was not meaningfully transported by SULP-4 [13]. However, it is possible that *sulp-4* mutations may enhance the formation of additional metabolic stones given a specific genetic or environmental perturbation. This hypothesis remains to be tested. Regardless, we think it is very intriguing that SLC26 homologs in mouse, humans, flies, and worms have all been shown to play roles in metabolic stone formation and suggest the potential for a fundamental mechanism underlying these discrete observations.

## Materials and methods

### General methods and strains

*C. elegans* were cultured using established protocols [22]. Briefly, animals were cultured at 20°C on nematode growth media (NGM) seeded with wild-type *E. coli* (OP50) unless otherwise noted. The wild-type strain of *C. elegans* was Bristol N2. Additional *E. coli* strains used in this work were BW25113 (Wild type, Moco+) and JW0764-2 (τι*moaA753::kan,* Moco-) [57].

*C. elegans* mutant and transgenic strains used in this work are listed here. When previously published, sources of strains are referenced. Unless a reference is provided, all strains were generated in this study.

#### Non-transgenic strains

N2, wild type [22]

GR2257, *cth-2(mg599) II* [8]

GR2259, *cth-2(mg599) II; moc-1(ok366) X* [8]

GR2260, *cdo-1(mg622)* [8]

GR2261, *cdo-1(mg622) moc-1(ok366) X* [8]

USD869, *xdh-1(ok3234) IV* (Outcrossed 2X)

USD1033, *sulp-4(rae319) V* (Outcrossed 4X)

USD1037, *sulp-4(rae319) V; cdo-1(mg622) moc-1(ok366) X*

USD1038, *sulp-4(rae319) V; cdo-1(mg622) X*

USD1055, *xdh-1(ok3234) IV; sulp-4(rae334) V*

USD1091, *sulp-4(rae334) V; cdo-1(mg622) X*

USD1103, *cth-2(mg599) II; sulp-4(rae334) V*

USD1105, *cth-2(mg599) II; xdh-1(ok3234) IV; sulp-4(rae334) V*

USD1146, *xdh-1(ok3234) IV; sulp-4(rae334) V; cdo-1(mg622) X*

USD1154, *xdh-1(ok3234) IV; cdo-1(mg622) X*

USD1163, *pnp-1(jy121) IV* (Outcrossed 1X) [25]

USD1170, *cth-2(mg599) II; xdh-1(ok3234) IV*

USD1174, *pnp-1(jy121) xdh-1(ok3234) IV*

USD1198, *pnp-1(jy121) IV; sulp-4(rae319) V*

USD1215, *pnp-1(jy121) xdh-1(ok3234) IV; sulp-4(rae319) V*

USD1230, *aex-5(sa23) I; xdh-1(ok3234) IV*

USD1269, *exc-5(rh232) xdh-1(ok3234) IV*

USD1308, *osm-8(n1518) II; xdh-1(ok3234) IV*

USD1310, *gpdh-1(ok1558) I* (Outcrossed 4X)

USD1312, *hmit-1.1(ok2923) V* (Outcrossed 4X)

USD1322, *gpdh-1(ok1558) I; sulp-4(rae319) V*

USD1324, *sulp-4(rae319) hmit-1.1(ok2923) V*

USD1327, *osm-8(n1518) II; xdh-1(ok3234) IV; cdo-1(mg622) X*

JT23, *aex-5(sa23) I* [30]

NJ731, *exc-5(rh232) IV* [28]

MT3571, *osm-8(n1518) II* [15]

RB1082, *sulp-5(ok1048) V*

RB1366, *sulp-2(ok1551) X*

RB1369, *sulp-2(ok1554) X*

RB1436, *sulp-1(ok1639) I*

RB2134, *sulp-8(ok2842) V*

FX08263, *sulp-5(tm8264) X*

VC3021, *sulp-7(ok3751) X*VC3045, *sulp-7(ok3752) X*

#### Transgenic strains

USD1060, *xdh-1(ok3234) IV; sulp-4(rae319) V; raeEx11*

USD1061, *xdh-1(ok3234) IV; sulp-4(rae319) V; raeEx119*

USD1062, *xdh-1(ok3234) IV; sulp-4(rae319) V; raeEx120*

USD1251, *qpIs11 I; sulp-4(rae319) V*

USD1277, *qpIs11 I; exc-5(rh232) IV*

BK36, *qpIs11 I; unc-119(ed3) III* [58]

#### EMS-derived strains

USD962*, *xdh-1(ok3234) IV; sulp-4(rae299) V*

USD1007, *xdh-1(ok3234) IV; sulp-4(rae299) V* (Outcrossed 2X)

USD971*, *xdh-1(ok3234) IV; sulp-4(rae302) V*

USD997, *xdh-1(ok3234) IV; sulp-4(rae302) V* (Outcrossed 1X)

USD990*, *xdh-1(ok3234) IV; sulp-4(rae319) V*

USD1001, *xdh-1(ok3234) IV; sulp-4(rae319) V* (Outcrossed 1X)

USD1013*, *xdh-1(ok3234) IV; sulp-4(rae320) V*

USD1019*, *xdh-1(ok3234) IV; sulp-4(rae326) V*

*Whole genome sequencing data for these *C. elegans* strains have been deposited at the NIH Sequence Read Archive (SRA) under accession PRJNA1208078.

#### CRIPSR/Cas9-derived stains

USD1042, *sulp-4(rae334) V*

### Chemical mutagenesis and whole genome sequencing

To define *C. elegans* gene activities that were necessary for promoting purine homeostasis, we carried out a chemical mutagenesis screen for mutations that enhanced the penetrance of xanthine stone formation in *xdh-1(ok3234)* mutant *C. elegans* (USD869). *C. elegans* were mutagenized with ethyl methanesulfonate (EMS) using established protocols [22]. Over multiple rounds of mutagenesis, we surveyed ∼100,000 mutagenized haploid genomes. To increase the likelihood of identifying a mutation that enhanced xanthine stone formation, we cloned ∼600 F2 generation animals that displayed a xanthine stone onto individual NGM petri dishes. F3 generation animals from these ∼600 isolates were then screened qualitatively for a population-level increase in xanthine stone penetrance. We demanded that new mutant strains of interest were viable and fertile.

Here we report the analysis of 5 new mutant strains (USD962, USD971, USD990, USD1013, and USD1019). Each of these strains carried new EMS-induced lesions (*rae299, rae302, rae319, rae320,* or *rae326)* that enhanced the formation of xanthine stones in an *xdh-1* mutant background. Each mutation was recessive; when heterozygous, each lesion caused 5% (*rae302, n*=42 individuals), 0% (*rae319, n*=38 individuals), 2% (*rae299, n*=41 individuals), or 5% (*rae326, n*=22 individuals) xanthine stone formation in an *xdh-1* mutant background, dramatically reduced when compared to their homozygous counterparts (**Fig. 2B)**. *rae320* was never characterized as dominant or recessive.

To further genetically analyze these lesions, we performed complementation analyses of these new mutations. The *rae319* lesion failed to complement *rae302* (100% xanthine stone penetrance, *n*=71 individuals), *rae299* (87% xanthine stone penetrance, *n*=38 individuals), and *rae326* (100% xanthine stone penetrance, *n*=14 individuals). All complementation experiments were performed in an *xdh-1(ok3234)* homozygous mutant genetic background. These results suggest *rae302, rae319, rae299,* and *rae326* all impact the same gene. Complementation studies were not performed on *rae320*.

To identify EMS-induced mutations in our strains of interest we followed established protocols [59]. Briefly, whole genomic DNA was prepared from *C. elegans* using the Gentra Puregene Tissue Kit (Qiagen) and genomic DNA libraries were prepared using the NEBNext genomic DNA library construction kit (New England Biolabs). DNA libraries were sequenced on an Illumina NovaSeq and deep sequencing reads were analyzed using standard methods on Galaxy, a web-based platform for computational analyses [60]. Briefly, sequencing reads were trimmed and aligned to the WBcel235 *C. elegans* reference genome [61, 62]. Variations from the reference genome and the putative impact of those variations were annotated and extracted for analysis [63–65]. All 4 strains that formed a complementation group possessed novel mutations in the gene *sulp-4,* strongly suggesting that these lesions in *sulp-4* caused the enhanced xanthine stone formation in the *xdh-1* mutant background (**Fig. 2A, Table S1**). Although the *rae320* lesion found in USD1013 was not analyzed via complementation, whole genome sequence analyses identified a homozygous mutation in *sulp-4*. Thus, we assume that the lesion in *sulp-4* found in USD1013 is also causative for the enhanced xanthine stone formation in the *xdh-1* mutant background (**Table S1)**. In fact, *rae319* and *rae320* are identical genetic lesions. We know that strains carrying these genetic lesions are not siblings as they were derived from independent rounds of mutagenesis. Whole genome sequencing data for these *C. elegans* strains have been deposited at the NIH Sequence Read Archive (SRA) under accession PRJNA1208078.

### CRISPR/Cas9 genome editing

Genome engineering using CRISPR/Cas9 technology was performed using established techniques [23, 24]. Briefly, 2 guide RNAs were designed and synthesized (IDT, crRNA) that targeted the *sulp-4* locus (5’-agagttagctttgtacaacg-3’ and 5’-atagcacatgatacttccgt-3’). Cas9 (IDT) guide RNA ribonucleoprotein complexes were directly injected into the *C. elegans* germline [23]. Newly induced deletions were identified in the offspring of injected animals using a PCR-based screening approach. The DNA primers used to screen for new deletions were: 5’-gcagagaaactcagagcaacaa-3’ and 5’-gcttggtttggaaactttgg-3’. We were able to isolate and homozygoze *sulp-4(rae334),* a new deletion of *sulp-4* (**Fig. 2A**).

### *C. elegans* transgenesis

Transgenic *C. elegans* carrying extrachromosomal arrays were generated by micro-injecting the gonad of young adult *xdh-1(ok3234); sulp-4(rae319)* double mutant *C. elegans* with an injection mix consisting of the *Psulp-4::SULP-4::GFP* plasmid (20 ng/μl), the *Pmyo-2::mCherry* co-injection marker (2 ng/μl), and the KB+ ladder (78 ng/μl, New England Biolabs) [66]. Three independently derived transgenic strains carrying the extrachromosomal arrays *raeEx118, raeEx119,* or *raeEx120* were isolated and maintained by propagating individual animals based on expression of the fluorescent mCherry protein in the pharynx.

### Determination of xanthine stone penetrance

To determine the percentage of animals that developed a xanthine stone, we cultured wild-type, mutant, and transgenic *C. elegans* beginning at the L1 stage of development under various growth conditions. Animals were assessed for the formation of a xanthine stone beginning at the L4 stage.

Animals were assessed daily through the first 4 days of adulthood except for experiments that used the *cdo-1(mg622)* allele or supplemental cysteine where assays were terminated at day 3 or 2 of adulthood respectively. *cdo-1(mg622)* and supplemental cysteine caused early lethality that limited the number of individuals that survived per biological replicate. Thus, the assays were shortened to increase the sample size in **Fig. 1B, 5A, 5B, 6C, and S4**. Importantly, all datapoints in a given figure panel were subjected to the same assay conditions and are thus directly comparable. Xanthine stones were determined based upon presence of exceptionally bright autofluorescent puncta that were opaque when observed by brightfield microscopy. If an individual displayed a stone, it was scored as such and removed from the assay. If an animal did not display a stone, it was counted and moved to a fresh petri dish to prevent contamination from the subsequent generation and allow for assessment on the following day. Thus, if an animal displayed a xanthine stone on any day of the assay, it scored positive and counted towards the penetrance of the phenotype. Xanthine stone penetrance was the percentage of animals that displayed a stone over the course of the assay. If animals went missing or died before the end of the assay, they were not included in the final analyses.

### *C. elegans* larval development and embryonic viability assays

To assay developmental rates, *C. elegans* were synchronized at the first stage of larval development. To synchronize animals, embryos were harvested from gravid adult animals via treatment with a bleach and sodium hydroxide solution. Embryos were then incubated overnight in M9 solution causing them to hatch and arrest development at the L1 stage [67]. Synchronized L1 animals were cultured for 72 hours under standard conditions, and live animals were imaged as described below. Animal length was measured from tip of head to the end of the tail.

To determine the hatching rate of wild-type and mutant *C. elegans,* we performed synchronized egg lays using young adult animals. Embryos were then scored for hatching ∼24 hours after being laid.

### Microscopy

Low magnification bright field and fluorescence images (**Fig. 1C and S1**) were collected using a Nikon SMZ25 microscope equipped with a Hamamatsu Orca flash 4.0 digital camera using NIS-Elements software (Nikon). High magnification differential interference contrast (DIC) and GFP fluorescence images (**Fig. 2C, 4A,B, and S3B-D**) were collected using a Nikon NiE microscope equipped with a Hamamatsu Orca flash 4.0 digital camera using NIS-Elements software (Nikon). Xanthine stones were visualized and imaged using the EGFP BP (FITC/Cy2) HC Filter Set (Nikon). All images were processed and analyzed using ImageJ software (NIH). All imaging was performed on live animals paralyzed using sodium azide.

### Quantitative PCR (qPCR)

RNA was extracted from synchronized wild-type, *sulp-4(rae319),* and *osm-8(n1518)* young adult animals using Trizol Reagent per manufacturer’s instructions (Invitrogen). Prior to RNA extraction, live *C. elegans* samples were washed and subsequently incubated for one hour in buffer M9 to allow for removal of bacterial contamination. cDNA was then synthesized using the GoScript Reverse Transcriptase System following manufacturer’s instructions (Promega). qPCR was performed using a CFX96 Real-Time System (Bio-Rad) and SYBR Green Master Mix following manufacturer’s instructions (Applied Biosystems). Relative mRNA levels were calculated using the comparative C_T_ methods [68]. Forward and reverse amplification primers were *act-1,* 5’-ctcttgccccatcaaccatg-3’ and 5’-cttgcttggagatccacatc-3’; *cdo-1,* 5’-ttcgatgagagaaccggaaag-3’ and 5’-gccattcttagatcctctgtagtc-3’; *hmit-1.1,* 5’-ccattgaagaggtagaaatgc-3’ and 5’-tgtacttcattgtgttgtcc-3’; and *gpdh-1,* 5’-tgcagagattccaggaaaccagg-3’ and 5’-cccttttgtagcttgccacggag-3’.

### Elemental analyses

For elemental analyses (S, K, Na, Mn, Fe, Cu, Zn, and Mo) of wild-type or *sulp-4(rae319)* mutant *C. elegans,* 29.9 to 34.7 mg of tissue (approximately 20,000 freeze-dried young adult *C. elegans*) were digested in 200μl of concentrated HNO_3_ at 90°C for one hour. After digestion, samples were diluted to a working volume of 1.4ml and subsequently diluted an additional 5X (Mn, Fe, Cu, Zn, and Mo) or 200X (S, K, and Na) to produce final samples for elemental analysis. All dilutions were performed with 1% HNO_3_. Inductively coupled plasma mass spectroscopy (ICP-MS) analysis was performed using an Agilent 8900 triple quad equipped with an SPS autosampler. The system was operated at a radio frequency power of 1550 W, an argon plasma gas flow rate of 15 L/min, and an Ar carrier gas flow rate of 0.9 L/min. Data were quantified using weighed, serial dilutions of a multi-element standard (CEM 2, (VHG Labs, VHG-SM70B-100) K, Na, Mn, Fe, Cu, Zn) and a single element standards for S (Spex CertiPrep, PLS9-2M) and Mo (Spex CertiPrep, CLMO9-2Y).

### NaCl tolerance assay

To assay NaCl tolerance, wild-type, *xdh-1, sulp-4,* or *osm-8* mutant *C. elegans* were synchronized at the first stage of larval development. To synchronize animals, embryos were harvested from gravid adult animals via treatment with a bleach and sodium hydroxide solution. Embryos were then incubated overnight in M9 solution causing them to hatch and arrest development at the L1 stage [67]. Synchronized L1 animals were cultured under standard conditions for 72 hours or until reaching the young adult stage. Young adult animals were then transferred to NGM plates with various NaCl concentrations (50, 200, 300, 400, 500, or 600mM NaCl). Animals were scored as alive or dead based on touch responsiveness after a 24-hour exposure to the various NaCl conditions. If animals went missing before the end of the assay, they were not included in the final analyses.

## Supporting information

Table S2, too large to embed in manuscript.

## Acknowledgements

Some *C. elegans* strains were provided by the CGC, which is funded by the NIH Office of Research Infrastructure Programs (P40 OD010440). We thank the lab of Emily Troemel for providing a *C. elegans* strain carrying the *pnp-1(jy121)* mutation. We thank the lab of Keith Nehrke for providing the *Psulp-4::SULP-4::GFP* plasmid (pTS1). Research reported in this publication was supported by the National Institute of General Medical Sciences of the National Institutes of Health under award number R35GM146871 (to K.W.). A.V.A. was supported by the National Science Foundation Division of Biological Infrastructure under award number 1756912. C.B. was supported by the National Institute of Childhood Health and Human Development of the National Institutes of Health under award number R25HD097633. ICP-MS measurements were performed in the OHSU Elemental Analysis Core with partial support from NIH (S10OD028492).

## Supporting Information

**Figure S1:**
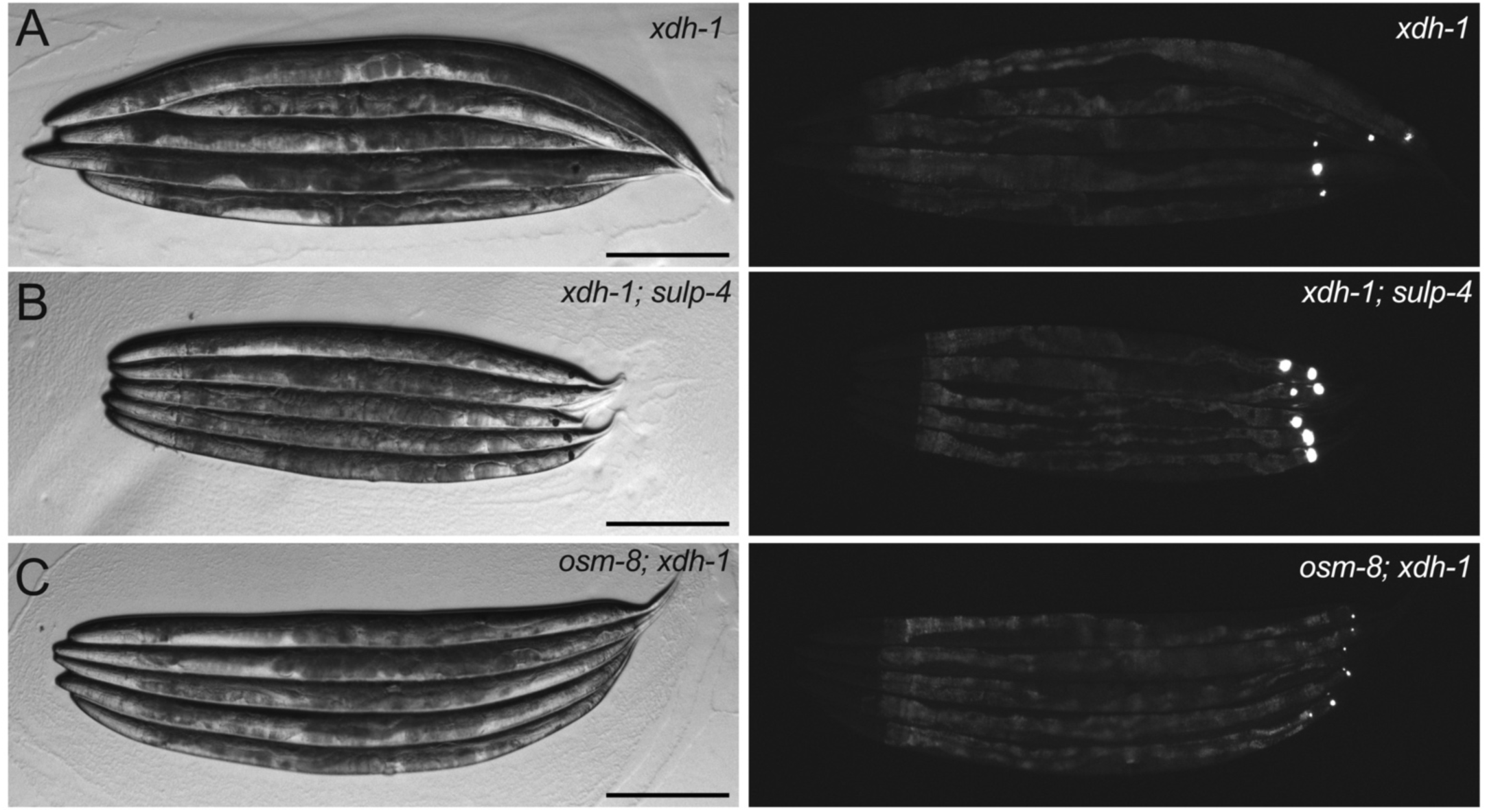
Loss of *sulp-4* enhanced the expressivity of the xanthine stone phenotype displayed by *xdh-1* mutant animals. Brightfield (left) and fluorescent (right) images of A) *xdh-1(ok3234),* B) *xdh-1(ok3234); sulp-4(rae319),* or C) *osm-8(n1518); xdh-1(ok3234)* mutant adult *C. elegans* cultured on wild-type *E. coli.* Scale bar is 250μm. Note, *xdh-1(ok3234)* animals displayed are older than their *xdh-1(ok3234); sulp-4(rae319)* or *osm-8(n1518); xdh-1(ok3234)* counterparts which were imaged as day 2 adults. This was necessary to allow us to identify sufficient *xdh-1* single mutant animals displaying a xanthine stone.

**Figure S2:**
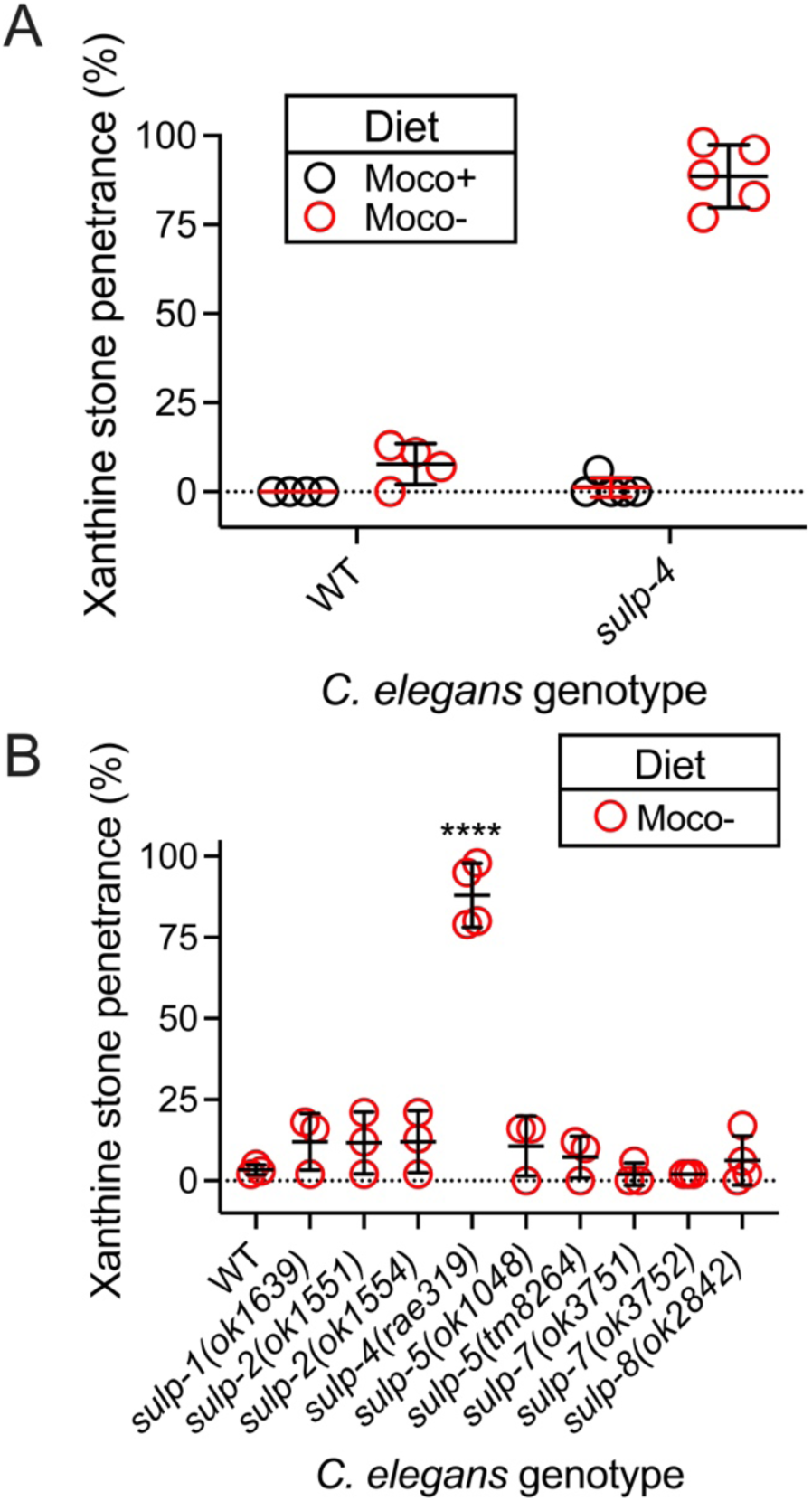
Inactivating mutations in *sulp* genes other than *sulp-4* did not enhance xanthine stone formation during dietary Moco deficiency. A) Wild-type and *sulp-4(rae319) C. elegans* were cultured on wild-type (black, Moco+) or *τιmoaA* mutant (red, Moco-) *E. coli* and assessed for the formation of xanthine stones. Note, data points for the wild-type animals cultured on Moco+ and Moco-*E. coli* are derived from the same experiment that is displayed in Fig. 1B However, in this analysis the animals were scored until day 4 of adulthood. B) Wild type and viable *sulp* mutant *C. elegans* were cultured on *τιmoaA* mutant (red, Moco-) *E. coli* and assessed for the formation of xanthine stones. Data points represent biological replicates. ****, p<0.0001, ordinary one-way ANOVA. If not indicated, data were not statistically different when compared to the wild type. Complete information regarding individuals scored per biological replicate are found in **Table S2**.

**Figure S3:**
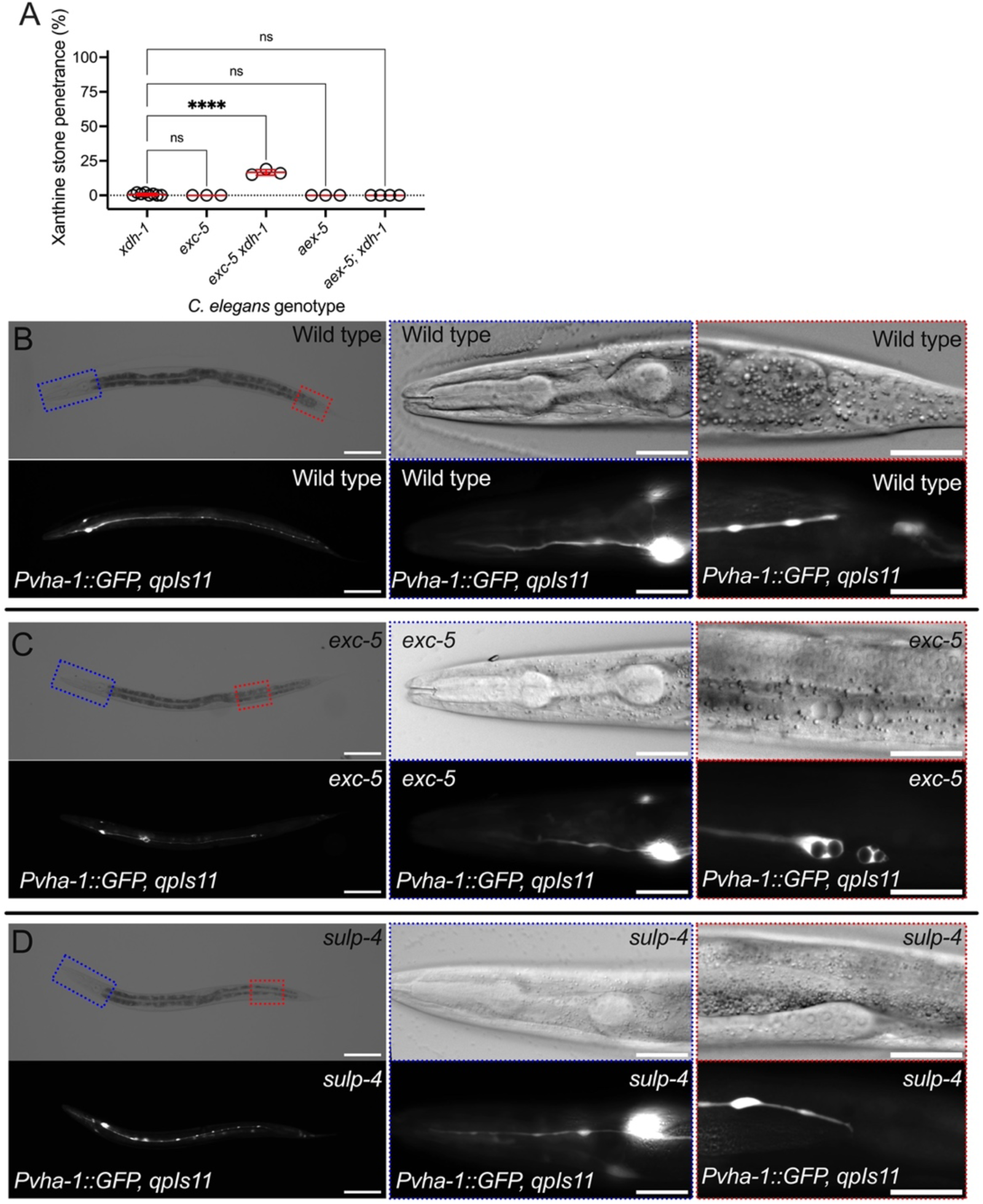
Loss of *exc-5* modestly enhanced the formation of xanthine stones in an *xdh-1* mutant background. A) *xdh-1(ok3234), exc-5(rh232), exc-5(rh232) xdh-1(ok3234), aex-5(sa23),* and *aex-5(sa23); xdh-1(ok3234)* mutant *C. elegans* were assessed for xanthine stone formation when cultured on wild-type *E. coli*. Individual data points represent biological replicates. Mean and standard deviation are displayed. ****, p<0.0001, ns, p>0.05, ordinary one-way ANOVA. Complete information regarding sample size and individuals scored per biological replicate are found in **Table S2**. B-D) Differential interference contrast (top) and fluorescence imaging (bottom) are displayed for B) wild-type, C) *exc-5(rh232),* or D) *sulp-4(rae319) C. elegans* expressing the *qpIs11* (*Pvha-1::GFP)* transgene, which marks the excretory cell. For images of whole animals (left panels), the blue (anterior) and red (posterior) boxes identify the regions displayed in the panels on the right. For images of whole animals (left panels), scale bar is 100μm. For images of anterior (blue box) and posterior (red box) excretory cell tubule extension, scale bar is 25μm.

**Figure S4:**
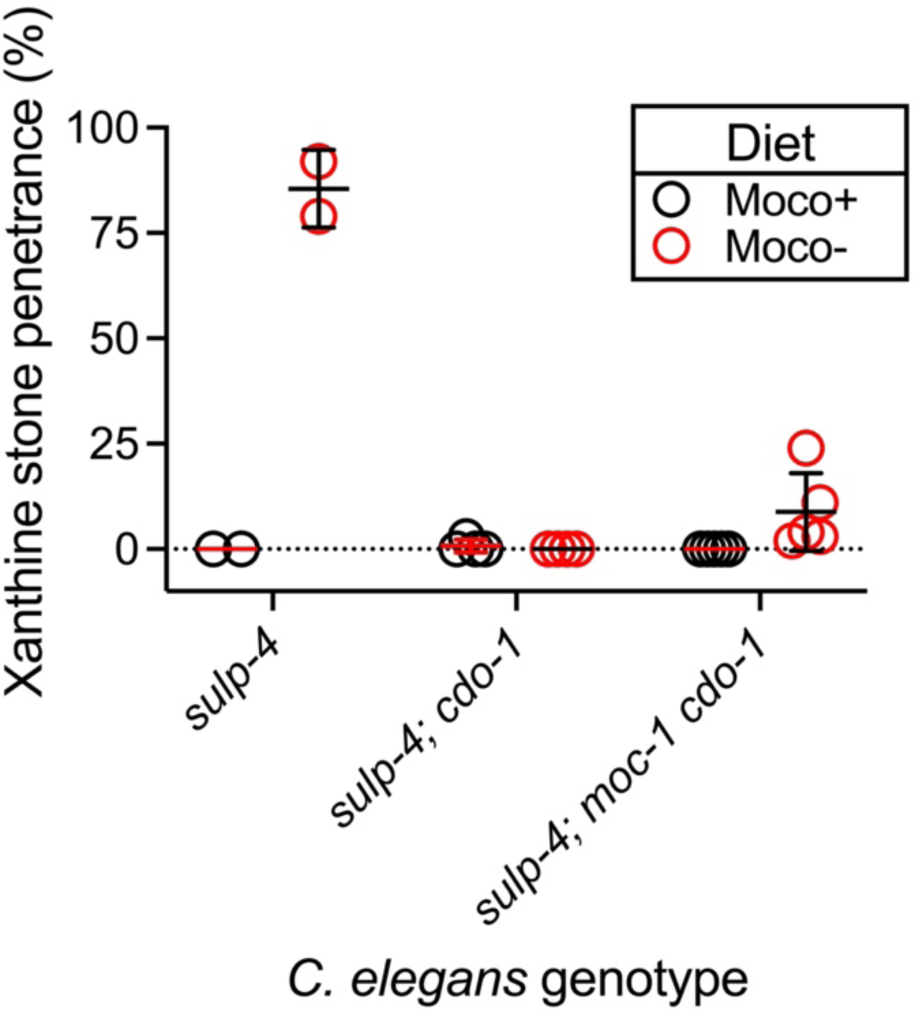
*cdo-1* was necessary for the enhanced formation of xanthine stones caused by *sulp-4* inactivation during dietary Moco deficiency. *sulp-4(rae319), sulp-4(rae319); cdo-1(mg622),* and *sulp-4(rae319); moc-1(ok366) cdo-1(mg622)* mutant *C. elegans* were cultured on wild-type (black, Moco+) or *τιmoaA* mutant (red, Moco-) *E. coli* and assessed for the formation of xanthine stones over the first 3 days of adulthood. Mean and standard deviation are displayed. Complete information regarding sample size and individuals scored per biological replicate are found in **Table S2**.

**Figure S5:**
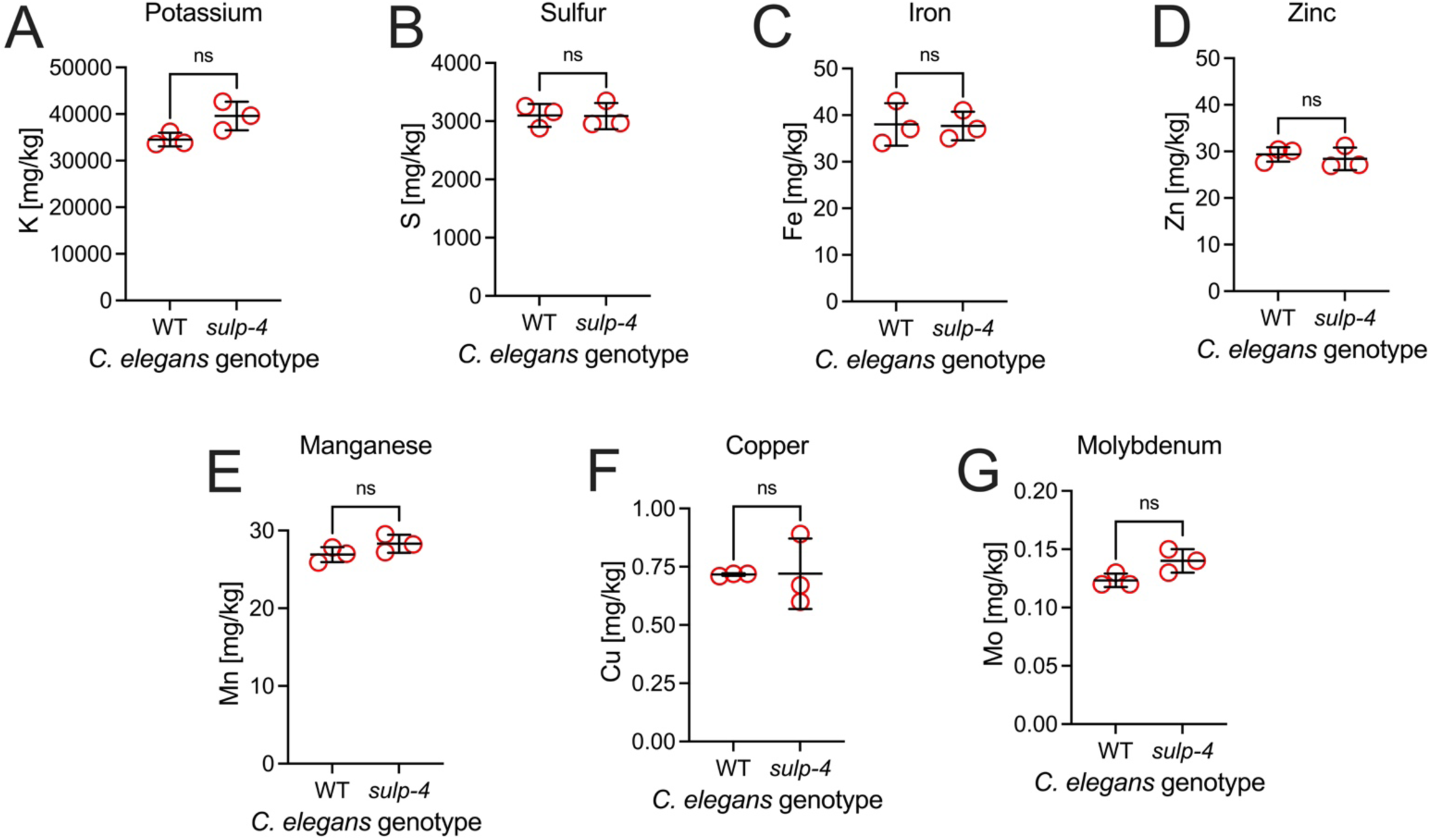
Elemental analyses of wild-type and *sulp-4* mutant *C. elegans*. ICP-MS analysis was performed on extracts from large cultures of young adult wild-type and *sulp-4(rae319)* mutant *C. elegans* to determine the concentrations of A) potassium, B) sulfur, C) iron, D) zinc, E) manganese, F) copper, and G) molybdenum. Individual data points represent biological replicates. Mean and standard deviation are displayed. ns, p>0.05, t test.

**Figure S6:**
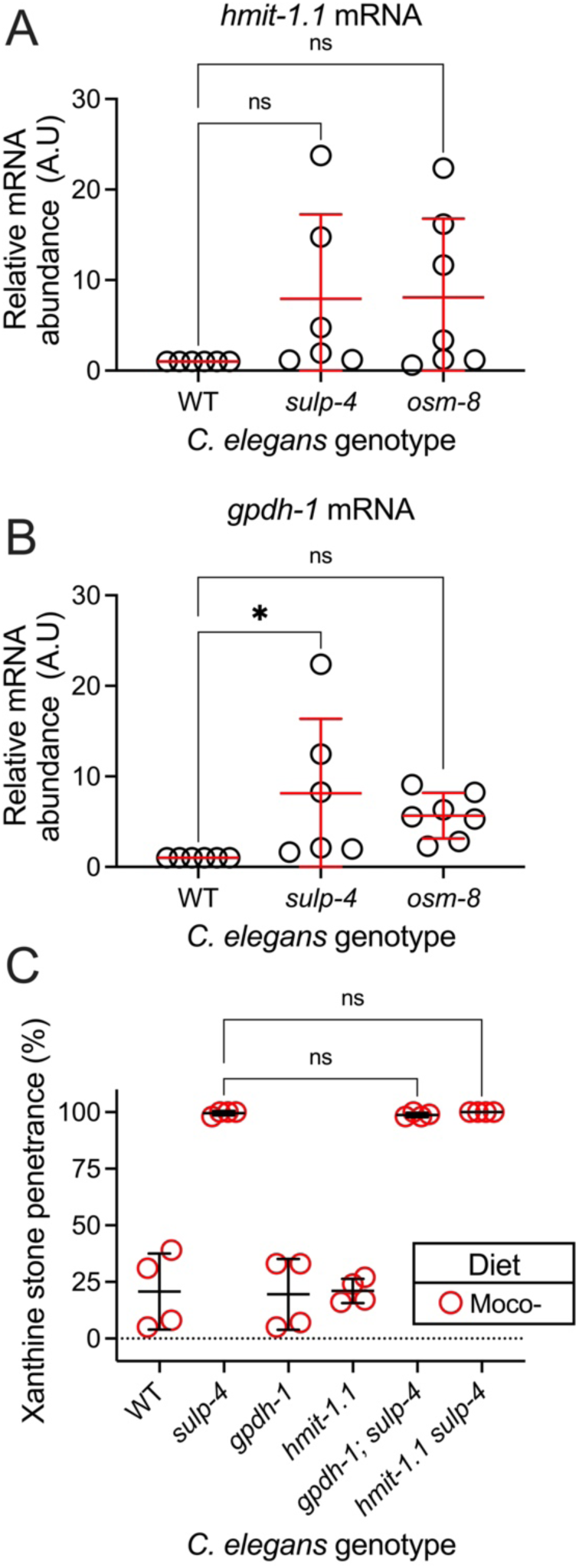
*hmit-1.1* and *gpdh-1* were not necessary for the enhanced formation of xanthine stones caused by *sulp-4* inactivation during dietary Moco deficiency. Relative mRNA expression of A) *hmit-1.1* and B) *gpdh-1* are displayed for total RNA isolated from wild-type, *sulp-4(rae319)*, and *osm-8(n1518)* young adult *C. elegans.* Relative mRNA abundance was determined via the delta-delta C_T_ method. All transcripts are normalized to *act-1.* Relative mRNA abundance for each transcript was set to one in the wild type. *, p<0.05, ns, p>0.05, ordinary one-way ANOVA. C) Wild type and mutant *C. elegans* were cultured on *τιmoaA* mutant (red, Moco-) *E. coli* and assessed for the formation of xanthine stones. Individual data points represent biological replicates. Mean and standard deviation are displayed. ns, p>0.05, ordinary one-way ANOVA. Complete information regarding sample size and individuals scored per biological replicate are found in **Table S2**.

**Table S1:** EMS-induced lesions in *sulp-4* that promoted the formation of xanthine stones in *xdh-1(ok3234)-*mutant *C. elegans*.

| Strain | Allele | Chromosome | Chromosome position | Gene affected | Reference base | Mutant base | Allele status | Variant impact |
| --- | --- | --- | --- | --- | --- | --- | --- | --- |
| USD962 | *rae299* | V | 11876913 | *sulp-4* | C | T | Homozygous | Pro325Leu |
| USD971 | *rae302* | V | 11877161 | *sulp-4* | G | A | Homozygous | Splice donor variant |
| USD990 | *rae319* | V | 11875799 | *sulp-4* | G | A | Homozygous | Gly116Glu |
| USD1013 | *rae320* | V | 11875799 | *sulp-4* | G | A | Homozygous | Gly116Glu |
| USD1019 | *rae326* | V | 11878726 | *sulp-4* | G | A | Homozygous | Splice acceptor variant |

**Table S2:** Raw data and information regarding sample sizes and biological replicates. Too large to be embedded in manuscript.

## References

1. Mullen NJ, Singh PK. Nucleotide metabolism: a pan-cancer metabolic dependency. Nat Rev Cancer. 2023;23(5):275–94. Epub 2023/03/28. doi: 10.1038/s41568-023-00557-7. PubMed PMID: 36973407; PubMed Central PMCID: PMCPMC10041518.

2. Lesch M, Nyhan WL. A FAMILIAL DISORDER OF URIC ACID METABOLISM AND CENTRAL NERVOUS SYSTEM FUNCTION. Am J Med. 1964;36:561–70. Epub 1964/04/01. doi: 10.1016/0002-9343(64)90104-4. PubMed PMID: 14142409.

3. Giblett ER, Ammann AJ, Wara DW, Sandman R, Diamond LK. Nucleoside-phosphorylase deficiency in a child with severely defective T-cell immunity and normal B-cell immunity. Lancet. 1975;1(7914):1010–3. Epub 1975/05/03. doi: 10.1016/s0140-6736(75)91950-9. PubMed PMID: 48676.

4. Duran M, Beemer FA, van de Heiden C, Korteland J, de Bree PK, Brink M, et al. Combined deficiency of xanthine oxidase and sulphite oxidase: a defect of molybdenum metabolism or transport? J Inherit Metab Dis. 1978;1(4):175–8. Epub 1978/01/01. doi: 10.1007/bf01805591. PubMed PMID: 117254.

5. Dent CE, Philpot GR. Xanthinuria, an inborn error (or deviation) of metabolism. Lancet. 1954;266(6804):182–5. Epub 1954/01/23. doi: 10.1016/s0140-6736(54)91257-x. PubMed PMID: 13118765.

6. Ball EG. XANTHINE OXIDASE: PURIFICATION AND PROPERTIES. J Biol Chem. 1939;128(1):51–67.

7. Sebesta I. Genetic disorders resulting in hyper- or hypouricemia. Adv Chronic Kidney Dis. 2012;19(6):398–403. Epub 2012/10/24. doi: 10.1053/j.ackd.2012.06.002. PubMed PMID: 23089275.

8. Warnhoff K, Ruvkun G. Molybdenum cofactor transfer from bacteria to nematode mediates sulfite detoxification. Nat Chem Biol. 2019;15(5):480–8. Epub 2019/03/27. doi: 10.1038/s41589-019-0249-y. PubMed PMID: 30911177; PubMed Central PMCID: PMCPMC6470025.

9. Schwarz G, Mendel RR, Ribbe MW. Molybdenum cofactors, enzymes and pathways. Nature. 2009;460(7257):839–47. Epub 2009/08/14. doi:10.1038/nature08302. PubMed PMID: 19675644.

10. Snoozy J, Breen PC, Ruvkun G, Warnhoff K. moc-6/MOCS2A is necessary for molybdenum cofactor synthesis in C. elegans. MicroPubl Biol. 2022;2022. Epub 2022/03/01. doi: 10.17912/micropub.biology.000531. PubMed PMID: 35224462; PubMed Central PMCID: PMCPMC8864482.

11. Warnhoff K, Hercher TW, Mendel RR, Ruvkun G. Protein-bound molybdenum cofactor is bioavailable and rescues molybdenum cofactor-deficient C. elegans. Genes Dev. 2021;35(3-4):212–7. Epub 2021/01/16. doi: 10.1101/gad.345579.120. PubMed PMID: 33446569; PubMed Central PMCID: PMCPMC7849362.

12. Oliphant KD, Fettig RR, Snoozy J, Mendel RR, Warnhoff K. Obtaining the necessary molybdenum cofactor for sulfite oxidase activity in the nematode Caenorhabditis elegans surprisingly involves a dietary source. J Biol Chem. 2023;299(1):102736. Epub 2022/11/25. doi: 10.1016/j.jbc.2022.102736. PubMed PMID: 36423681; PubMed Central PMCID: PMCPMC9793310.

13. Sherman T, Chernova MN, Clark JS, Jiang L, Alper SL, Nehrke K. The abts and sulp families of anion transporters from Caenorhabditis elegans. Am J Physiol Cell Physiol. 2005;289(2):C341–51. Epub 2005/04/09. doi: 10.1152/ajpcell.00071.2005. PubMed PMID: 15814591.

14. Nelson FK, Riddle DL. Functional study of the Caenorhabditis elegans secretory-excretory system using laser microsurgery. J Exp Zool. 1984;231(1):45–56. Epub 1984/07/01. doi: 10.1002/jez.1402310107. PubMed PMID: 6470649.

15. Rohlfing AK, Miteva Y, Moronetti L, He L, Lamitina T. The Caenorhabditis elegans mucin-like protein OSM-8 negatively regulates osmosensitive physiology via the transmembrane protein PTR-23. PLoS Genet. 2011;7(1):e1001267. Epub 2011/01/22. doi: 10.1371/journal.pgen.1001267. PubMed PMID: 21253570; PubMed Central PMCID: PMCPMC3017116.

16. Chalmers RA, Watts RW, Pallis C, Bitensky L, Chayen J. Crystalline deposits in striped muscle in xanthinuria. Nature. 1969;221(5176):170-1. Epub 1969/01/11. doi: 10.1038/221170a0. PubMed PMID: 5782709.

17. Watts RW, Engelman K, Klinenberg JR, Seegmiller JE, Sjoerdsma A. ENZYME DEFECT IN A CASE OF XANTHINURIA. Nature. 1964;201:395–6. Epub 1964/01/25. doi: 10.1038/201395a0. PubMed PMID: 14110004.

18. Ma X, Wang W, Bittner F, Schmidt N, Berkey R, Zhang L, et al. Dual and Opposing Roles of Xanthine Dehydrogenase in Defense-Associated Reactive Oxygen Species Metabolism in Arabidopsis. Plant Cell. 2016;28(5):1108–26. Epub 2016/05/07. doi: 10.1105/tpc.15.00880. PubMed PMID: 27152019; PubMed Central PMCID: PMCPMC4904670.

19. Chi T, Kim MS, Lang S, Bose N, Kahn A, Flechner L, et al. A Drosophila model identifies a critical role for zinc in mineralization for kidney stone disease. PLoS One. 2015;10(5):e0124150. Epub 2015/05/15. doi: 10.1371/journal.pone.0124150. PubMed PMID: 25970330; PubMed Central PMCID: PMCPMC4430225.

20. Glassman E, Mitchell HK. Mutants of Drosophila Melanogaster Deficient in Xanthine Dehydrogenase. Genetics. 1959;44(2):153–62. Epub 1959/03/01. doi: 10.1093/genetics/44.2.153. PubMed PMID: 17247815; PubMed Central PMCID: PMCPMC1209939.

21. Takagaki N, Ohta A, Ohnishi K, Kawanabe A, Minakuchi Y, Toyoda A, et al. The mechanoreceptor DEG-1 regulates cold tolerance in Caenorhabditis elegans. EMBO Rep. 2020;21(3):e48671. Epub 2020/02/06. doi: 10.15252/embr.201948671. PubMed PMID: 32009302; PubMed Central PMCID: PMCPMC7054665.

22. Brenner S. The genetics of Caenorhabditis elegans. Genetics. 1974;77(1):71–94. Epub 1974/05/01. PubMed PMID: 4366476; PubMed Central PMCID: PMCPMC1213120.

23. Cho SW, Lee J, Carroll D, Kim JS, Lee J. Heritable gene knockout in Caenorhabditis elegans by direct injection of Cas9-sgRNA ribonucleoproteins. Genetics. 2013;195(3):1177–80. Epub 2013/08/28. doi: 10.1534/genetics.113.155853. PubMed PMID: 23979576; PubMed Central PMCID: PMCPMC3813847.

24. Ghanta KS, Mello CC. Melting dsDNA Donor Molecules Greatly Improves Precision Genome Editing in Caenorhabditis elegans. Genetics. 2020;216(3):643–50. Epub 2020/09/24. doi: 10.1534/genetics.120.303564. PubMed PMID: 32963112; PubMed Central PMCID: PMCPMC7648581.

25. Tecle E, Chhan CB, Franklin L, Underwood RS, Hanna-Rose W, Troemel ER. The purine nucleoside phosphorylase pnp-1 regulates epithelial cell resistance to infection in C. elegans. PLoS Pathog. 2021;17(4):e1009350. Epub 2021/04/21. doi: 10.1371/journal.ppat.1009350. PubMed PMID: 33878133; PubMed Central PMCID: PMCPMC8087013.

26. Altschul SF, Gish W, Miller W, Myers EW, Lipman DJ. Basic local alignment search tool. J Mol Biol. 1990;215(3):403–10. Epub 1990/10/05. doi: 10.1016/s0022-2836(05)80360-2. PubMed PMID: 2231712.

27. Buechner M. Tubes and the single C. elegans excretory cell. Trends Cell Biol. 2002;12(10):479–84. Epub 2002/11/21. doi: 10.1016/s0962-8924(02)02364-4. PubMed PMID: 12441252.

28. Suzuki N, Buechner M, Nishiwaki K, Hall DH, Nakanishi H, Takai Y, et al. A putative GDP-GTP exchange factor is required for development of the excretory cell in Caenorhabditis elegans. EMBO Rep. 2001;2(6):530–5. Epub 2001/06/21. doi: 10.1093/embo-reports/kve110. PubMed PMID: 11415987; PubMed Central PMCID: PMCPMC1083904.

29. Branicky R, Hekimi S. What keeps C. elegans regular: the genetics of defecation. Trends Genet. 2006;22(10):571–9. Epub 2006/08/17. doi: 10.1016/j.tig.2006.08.006. PubMed PMID: 16911844.

30. Thomas JH. Genetic analysis of defecation in Caenorhabditis elegans. Genetics. 1990;124(4):855–72. Epub 1990/04/01. doi: 10.1093/genetics/124.4.855. PubMed PMID: 2323555; PubMed Central PMCID: PMCPMC1203977.

31. Warnhoff K, Bhattacharya S, Snoozy J, Breen PC, Ruvkun G. Hypoxia-inducible factor induces cysteine dioxygenase and promotes cysteine homeostasis in Caenorhabditis elegans. Elife. 2024;12. Epub 2024/02/13. doi: 10.7554/eLife.89173. PubMed PMID: 38349720.

32. Stipanuk MH, Ueki I. Dealing with methionine/homocysteine sulfur: cysteine metabolism to taurine and inorganic sulfur. J Inherit Metab Dis. 2011;34(1):17–32. Epub 2010/02/18. doi: 10.1007/s10545-009-9006-9. PubMed PMID: 20162368; PubMed Central PMCID: PMCPMC2901774.

33. Dissanayake LV, Zietara A, Levchenko V, Spires DR, Burgos Angulo M, El-Meanawy A, et al. Lack of xanthine dehydrogenase leads to a remarkable renal decline in a novel hypouricemic rat model. iScience. 2022;25(9):104887. Epub 2022/08/31. doi: 10.1016/j.isci.2022.104887. PubMed PMID: 36039296; PubMed Central PMCID: PMCPMC9418856.

34. Rohlfing AK, Miteva Y, Hannenhalli S, Lamitina T. Genetic and physiological activation of osmosensitive gene expression mimics transcriptional signatures of pathogen infection in C. elegans. PLoS One. 2010;5(2):e9010. Epub 2010/02/04. doi: 10.1371/journal.pone.0009010. PubMed PMID: 20126308; PubMed Central PMCID: PMCPMC2814864.

35. Lamitina T, Huang CG, Strange K. Genome-wide RNAi screening identifies protein damage as a regulator of osmoprotective gene expression. Proc Natl Acad Sci U S A. 2006;103(32):12173–8. Epub 2006/08/02. doi: 10.1073/pnas.0602987103. PubMed PMID: 16880390; PubMed Central PMCID: PMCPMC1567714.

36. Lamitina ST, Morrison R, Moeckel GW, Strange K. Adaptation of the nematode Caenorhabditis elegans to extreme osmotic stress. Am J Physiol Cell Physiol. 2004;286(4):C785–91. Epub 2003/12/03. doi: 10.1152/ajpcell.00381.2003. PubMed PMID: 14644776.

37. Kage-Nakadai E, Uehara T, Mitani S. H+/myo-inositol transporter genes, hmit-1.1 and hmit-1.2, have roles in the osmoprotective response in Caenorhabditis elegans. Biochem Biophys Res Commun. 2011;410(3):471–7. Epub 2011/06/18. doi: 10.1016/j.bbrc.2011.06.001. PubMed PMID: 21679696.

38. Lee SD, Choi SY, Lim SW, Lamitina ST, Ho SN, Go WY, et al. TonEBP stimulates multiple cellular pathways for adaptation to hypertonic stress: organic osmolyte-dependent and - independent pathways. Am J Physiol Renal Physiol. 2011;300(3):F707–15. Epub 2011/01/07. doi: 10.1152/ajprenal.00227.2010. PubMed PMID: 21209002; PubMed Central PMCID: PMCPMC3064130.

39. Pais VM, Jr., Lowe G, Lallas CD, Preminger GM, Assimos DG. Xanthine urolithiasis. Urology. 2006;67(5):1084.e9–11. Epub 2006/05/16. doi: 10.1016/j.urology.2005.10.057. PubMed PMID: 16698380.

40. Yanagi KS, Jochim B, Kunjo SO, Breen P, Ruvkun G, Lehrbach N. Mutations in nucleotide metabolism genes bypass proteasome defects in png-1/NGLY1-deficient Caenorhabditis elegans. PLoS Biol. 2024;22(7):e3002720. Epub 2024/07/11. doi: 10.1371/journal.pbio.3002720. PubMed PMID: 38991033; PubMed Central PMCID: PMCPMC11265709.

41. Ast T, Meisel JD, Patra S, Wang H, Grange RMH, Kim SH, et al. Hypoxia Rescues Frataxin Loss by Restoring Iron Sulfur Cluster Biogenesis. Cell. 2019;177(6):1507–21.e16. Epub 2019/04/30. doi: 10.1016/j.cell.2019.03.045. PubMed PMID: 31031004; PubMed Central PMCID: PMCPMC6911770.

42. Bantia S, Parker C, Upshaw R, Cunningham A, Kotian P, Kilpatrick JM, et al. Potent orally bioavailable purine nucleoside phosphorylase inhibitor BCX-4208 induces apoptosis in B- and T-lymphocytes--a novel treatment approach for autoimmune diseases, organ transplantation and hematologic malignancies. Int Immunopharmacol. 2010;10(7):784–90. Epub 2010/04/20. doi: 10.1016/j.intimp.2010.04.009. PubMed PMID: 20399911.

43. Kamath VP, Juarez-Brambila JJ, Morris PE, Jr. Synthesis of labeled BCX-4208, a potent inhibitor of purine nucleoside phosphorylase. Drug Test Anal. 2009;1(3):125–7. Epub 2010/04/01. doi: 10.1002/dta.24. PubMed PMID: 20355185.

44. Adrogué HJ, Madias NE. Hypernatremia. N Engl J Med. 2000;342(20):1493–9. Epub 2000/05/18. doi: 10.1056/nejm200005183422006. PubMed PMID: 10816188.

45. Prince CL, Scardino PL, Wolan CT. The effect of temperature, humidity and dehydration on the formation of renal calculi. J Urol. 1956;75(2):209–15. Epub 1956/02/01. doi: 10.1016/s0022-5347(17)66798-3. PubMed PMID: 13296117.

46. Mudd SH, Irreverre F, Laster L. Sulfite oxidase deficiency in man: demonstration of the enzymatic defect. Science. 1967;156(3782):1599–602. Epub 1967/06/23. doi: 10.1126/science.156.3782.1599. PubMed PMID: 6025118.

47. Dominy JE, Jr., Hirschberger LL, Coloso RM, Stipanuk MH. In vivo regulation of cysteine dioxygenase via the ubiquitin-26S proteasome system. Adv Exp Med Biol. 2006;583:37–47. Epub 2006/12/13. doi: 10.1007/978-0-387-33504-9_4. PubMed PMID: 17153587.

48. Dominy JE, Jr., Hirschberger LL, Coloso RM, Stipanuk MH. Regulation of cysteine dioxygenase degradation is mediated by intracellular cysteine levels and the ubiquitin-26 S proteasome system in the living rat. Biochem J. 2006;394(Pt 1):267–73. Epub 2005/11/03. doi: 10.1042/bj20051510. PubMed PMID: 16262602; PubMed Central PMCID: PMCPMC1386025.

49. Stipanuk MH, Hirschberger LL, Londono MP, Cresenzi CL, Yu AF. The ubiquitin-proteasome system is responsible for cysteine-responsive regulation of cysteine dioxygenase concentration in liver. Am J Physiol Endocrinol Metab. 2004;286(3):E439–48. Epub 2003/12/03. doi: 10.1152/ajpendo.00336.2003. PubMed PMID: 14644768.

50. Lee JI, Londono M, Hirschberger LL, Stipanuk MH. Regulation of cysteine dioxygenase and gamma-glutamylcysteine synthetase is associated with hepatic cysteine level. J Nutr Biochem. 2004;15(2):112–22. Epub 2004/02/20. doi: 10.1016/j.jnutbio.2003.10.005. PubMed PMID: 14972351.

51. Kwon YH, Stipanuk MH. Cysteine regulates expression of cysteine dioxygenase and gamma-glutamylcysteine synthetase in cultured rat hepatocytes. Am J Physiol Endocrinol Metab. 2001;280(5):E804–15. Epub 2001/04/05. doi: 10.1152/ajpendo.2001.280.5.E804. PubMed PMID: 11287364.

52. Hill AJ, Basourakos SP, Lewicki P, Wu X, Arenas-Gallo C, Chuang D, et al. Incidence of Kidney Stones in the United States: The Continuous National Health and Nutrition Examination Survey. J Urol. 2022;207(4):851–6. Epub 2021/12/03. doi: 10.1097/ju.0000000000002331. PubMed PMID: 34854755.

53. Jiang Z, Asplin JR, Evan AP, Rajendran VM, Velazquez H, Nottoli TP, et al. Calcium oxalate urolithiasis in mice lacking anion transporter Slc26a6. Nat Genet. 2006;38(4):474–8. Epub 2006/03/15. doi: 10.1038/ng1762. PubMed PMID: 16532010.

54. Cornière N, Thomson RB, Thauvin S, Villoutreix BO, Karp S, Dynia DW, et al. Dominant negative mutation in oxalate transporter SLC26A6 associated with enteric hyperoxaluria and nephrolithiasis. J Med Genet. 2022;59(11):1035–43. Epub 2022/02/05. doi: 10.1136/jmedgenet-2021-108256. PubMed PMID: 35115415; PubMed Central PMCID: PMCPMC9346097.

55. Gee HY, Jun I, Braun DA, Lawson JA, Halbritter J, Shril S, et al. Mutations in SLC26A1 Cause Nephrolithiasis. Am J Hum Genet. 2016;98(6):1228–34. Epub 2016/05/24. doi: 10.1016/j.ajhg.2016.03.026. PubMed PMID: 27210743; PubMed Central PMCID: PMCPMC4908148.

56. Hirata T, Cabrero P, Berkholz DS, Bondeson DP, Ritman EL, Thompson JR, et al. In vivo Drosophilia genetic model for calcium oxalate nephrolithiasis. Am J Physiol Renal Physiol. 2012;303(11):F1555–62. Epub 2012/09/21. doi: 10.1152/ajprenal.00074.2012. PubMed PMID: 22993075; PubMed Central PMCID: PMCPMC3532482.

57. Baba T, Ara T, Hasegawa M, Takai Y, Okumura Y, Baba M, et al. Construction of Escherichia coli K-12 in-frame, single-gene knockout mutants: the Keio collection. Mol Syst Biol. 2006;2:2006.0008. Epub 2006/06/02. doi: 10.1038/msb4100050. PubMed PMID: 16738554; PubMed Central PMCID: PMCPMC1681482.

58. Mattingly BC, Buechner M. The FGD homologue EXC-5 regulates apical trafficking in C. elegans tubules. Dev Biol. 2011;359(1):59–72. Epub 2011/09/06. doi: 10.1016/j.ydbio.2011.08.011. PubMed PMID: 21889936; PubMed Central PMCID: PMCPMC3212395.

59. Lehrbach NJ, Ji F, Sadreyev R. Next-Generation Sequencing for Identification of EMS-Induced Mutations in Caenorhabditis elegans. Curr Protoc Mol Biol. 2017;117:7.29.1–7..12. Epub 2017/01/07. doi: 10.1002/cpmb.27. PubMed PMID: 28060408; PubMed Central PMCID: PMCPMC5303615.

60. Community G. The Galaxy platform for accessible, reproducible and collaborative biomedical analyses: 2022 update. Nucleic Acids Res. 2022;50(W1):W345–51. Epub 2022/04/22. doi: 10.1093/nar/gkac247. PubMed PMID: 35446428; PubMed Central PMCID: PMCPMC9252830.

61. Bolger AM, Lohse M, Usadel B. Trimmomatic: a flexible trimmer for Illumina sequence data. Bioinformatics. 2014;30(15):2114–20. Epub 2014/04/04. doi: 10.1093/bioinformatics/btu170. PubMed PMID: 24695404; PubMed Central PMCID: PMCPMC4103590.

62. Li H, Durbin R. Fast and accurate long-read alignment with Burrows-Wheeler transform. Bioinformatics. 2010;26(5):589–95. Epub 2010/01/19. doi: 10.1093/bioinformatics/btp698. PubMed PMID: 20080505; PubMed Central PMCID: PMCPMC2828108.

63. Wilm A, Aw PP, Bertrand D, Yeo GH, Ong SH, Wong CH, et al. LoFreq: a sequence-quality aware, ultra-sensitive variant caller for uncovering cell-population heterogeneity from high-throughput sequencing datasets. Nucleic Acids Res. 2012;40(22):11189–201. Epub 2012/10/16. doi: 10.1093/nar/gks918. PubMed PMID: 23066108; PubMed Central PMCID: PMCPMC3526318.

64. Cingolani P, Patel VM, Coon M, Nguyen T, Land SJ, Ruden DM, et al. Using Drosophila melanogaster as a Model for Genotoxic Chemical Mutational Studies with a New Program, SnpSift. Front Genet. 2012;3:35. Epub 2012/03/22. doi: 10.3389/fgene.2012.00035. PubMed PMID: 22435069; PubMed Central PMCID: PMCPMC3304048.

65. Cingolani P, Platts A, Wang le L, Coon M, Nguyen T, Wang L, et al. A program for annotating and predicting the effects of single nucleotide polymorphisms, SnpEff: SNPs in the genome of Drosophila melanogaster strain w1118; iso-2; iso-3. Fly (Austin). 2012;6(2):80–92. Epub 2012/06/26. doi: 10.4161/fly.19695. PubMed PMID: 22728672; PubMed Central PMCID: PMCPMC3679285.

66. Mello CC, Kramer JM, Stinchcomb D, Ambros V. Efficient gene transfer in C.elegans: extrachromosomal maintenance and integration of transforming sequences. Embo j. 1991;10(12):3959–70. Epub 1991/12/01. doi: 10.1002/j.1460-2075.1991.tb04966.x. PubMed PMID: 1935914; PubMed Central PMCID: PMCPMC453137.

67. Stiernagle T. Maintenance of C. elegans. WormBook. 2006:1–11. Epub 2007/12/01. doi: 10.1895/wormbook.1.101.1. PubMed PMID: 18050451; PubMed Central PMCID: PMCPMC4781397.

68. Schmittgen TD, Livak KJ. Analyzing real-time PCR data by the comparative C(T) method. Nat Protoc. 2008;3(6):1101–8. Epub 2008/06/13. doi: 10.1038/nprot.2008.73. PubMed PMID: 18546601.

